# Voltage-Seq: all-optical postsynaptic connectome-guided single-cell transcriptomics

**DOI:** 10.1101/2022.11.09.515875

**Authors:** Veronika Csillag, Marianne Hiriart Bizzozzero, Joyce Noble, Björn Reinius, János Fuzik

## Abstract

Understanding the routing of neuronal information requires the functional characterization of connections. Neuronal projections recruit large postsynaptic ensembles with distinct postsynaptic response types (PRTs). PRT is probed by low-throughput whole-cell electrophysiology and is not a selection criterion for single-cell RNA-sequencing (scRNA-seq). To overcome these limitations and target neurons based on specific PRTs for soma harvesting and subsequent scRNA-seq we created Voltage-Seq. To test our methodology, we established all-optical voltage imaging and recorded the PRT of 8347 periaqueductal gray (PAG) neurons evoked by the optogenetic activation of ventromedial hypothalamic (VMH) terminals. PRTs were classified and spatially resolved in the entire VMH-PAG connectome. We built an on-site analysis named VoltView to navigate soma harvesting towards target PRTs guided by a classifier which used the VMH-PAG connectome database as a reference. We demonstrated the agility of Voltage-Seq in locating VMH-driven GABAergic neurons in the PAG, solely guided by the on-site classification in VoltView.

## Main

Neuronal information flows through synaptic connections and modulates large postsynaptic populations in a cell-type-specific manner. Neuronal types can be characterized by morphology, anatomic position, intrinsic excitability, gene expression profile and connectivity. Patch-Seq^1–3^ pioneered the molecular characterization of neurons classified by whole-cell patch-clamp^4^ recording of intrinsic excitability. To date, synaptic connectivity is probed with whole-cell patchclamp as it requires the detection of subthreshold postsynaptic potentials (PSPs). The throughput of this technique (~dozen neurons per day) is low for the efficient mapping of diverse PRTs in a large postsynaptic population. Finding neurons with specific PRTs for further investigation requires high-throughput connectivity testing and the detection of both subthreshold and suprathreshold membrane potential changes. Genetically encoded fluorescent voltage indicators (GEVIs)^5,6^ faithfully report subthreshold voltage changes of both polarities^5^ with reliable temporal dynamics and can capture single action potentials (APs). Voltage imaging has high throughput^7^ for simultaneous optical recording of dozens of neurons.

The PAG is a midbrain structure processing panicogenic stimuli^8^, and involved in the regulation of autonomic functions^9^ and motivated behaviors^10^. PAG receives a strong excitatory input from the VMH^11^. The VMH-PAG axons cover a 2-mm-long anterior-posterior (A-P) range of the dorsal, dorsolateral, and lateral (d, dl, l-PAG) portions of the PAG. The cell-type- and circuit-motif-specific routing of VMH information in the local PAG circuitry is poorly understood due to its large anatomical extension and high neuronal diversity^12^. We used the VMH-PAG pathway as a model to optimize the Voltage-Seq methodology, to all-optical voltage image PRTs, and to select specific neurons for somatic harvesting and subsequent scRNA-seq.

First, we set up all-optical voltage imaging *ex vivo* implementing the Voltron sensor^7^. Our tiled all-optical imaging has a high throughput of probing up to 1000-1500 connections per animal. Next, we generated a whole-structure synaptic connectome of the VMH-PAG projection. Spatial mapping of this connectome revealed the topography of the distinct PRTs in the entire PAG. Next, we built an interactive on-site analysis named VoltView which gave an overview of ~30-80 all-optical imaged PRTs in 1 minute. We added a classifier incorporating the generated VMH-PAG connectome data. With that, VoltView could on-site-classify PRTs and navigate a recording- or harvesting pipette to neurons with user-defined target PRTs. We tested Voltage-Seq to locate sparse GABAergic neurons in the VMH-PAG guided by the on-site analysis in VoltView. Remarkably, for the first time, we identified a d-dlPAG-located GABAergic feed-forward disinhibitory circuit motif driven by the glutamatergic VMH and using transcriptomics we identified a neuromodulator that regulates this disinhibitory motif.

## Results

### All-optical postsynaptic voltage imaging

We established and optimized all-optical voltage imaging *ex vivo*, using the recently introduced Voltron^7^ sensor. We co-injected a virus to express Cre recombinase and another to express Cre-dependent soma-targeting Voltron (Voltron-ST) (Fig. 1a). With this viral combination we achieved a cell-type-independent Voltron-ST labeling in ~35-40% of PAG neurons (Extended Data Fig. 1a). For VMH-PAG all-optical connectivity testing we also expressed Channelrhodopsin-2 (ChR2) in the VMH (Fig 1a). We fluorescent-labeled the Voltron with the Janelia Fluor^®^585-HaloTag (JF-585) and configured the light path accordingly (Extended Data Fig. 1b). We optimized acquisition speed to 600 Hz which captured all the optical action potentials (o-APs) in 3-4 frames (Extended Data Fig. 1c). At this framerate, firing rate up to ~125 Hz and changes in o-AP half-width could be detected (Extended Data Fig. 1d,e,f). With simulated PSPs we validated the detection limit of ~2-3 mV optical-excitatory, and inhibitory PSPs (o-EPSPs, o-IPSPs) (Extended Data Fig. 1g,h). We validated that an extreme ~21 mW JF-585 excitation had a negligible ~3 mV cross-activation of ChR2 (Extended Data Fig. 2a,b,c). We validated that ChR2-evoked synaptic release was not influenced by cross-activation of JF-585 excitation (Extended Data Fig. 2d). We successfully all-optical-imaged both VMH-PAG o-EPSPs and o-IPSPs confirmed by paralleled e-IPSP recording in the imaged PAG neuron (Fig. 1b). The 3-4 frame delay of o-IPSP and 4.7±0.4ms e-IPSP latency indicated the putative disynaptic nature of the response (Fig. 1b). All-optical recordings revealed 473 nm light-induced narrow artefacts with reversed polarity to o-APs, which were removed (Extended Data Fig. 2e). We could detect compound o-EPSP/o-IPSP responses with a narrow profile (Fig. 1c), which displayed o-IPSP upon depolarization (Extended Data Fig. 2f,). We used antagonist pharmacology to dissect a compound o-PSP of a putative disynaptic motif (Fig. 1c,d). A rebound-bursting neuron confirmed the co-detection of large slow hyperpolarization (300-500 ms), rhythmic depolarization and fast spiking (Fig. 1e). The involvement of γ-aminobutyric acid (GABA) was predicted by the compound o-PSPs during optical stimulation (Op). GABA antagonist pharmacology turned rebound-bursting into onset-bursting confirming the inhibitory component of the compound o-PSPs (Extended Data Fig. 2g). Substantial bleaching of JF-585 occurred after ~3-4 minutes of imaging (Extended Data Fig. 2h). Taken together, optimized all-optical imaging could detect the firing activity, bursting and mono- and disynaptic excitatory and inhibitory subthreshold events.

**Fig. 1:**
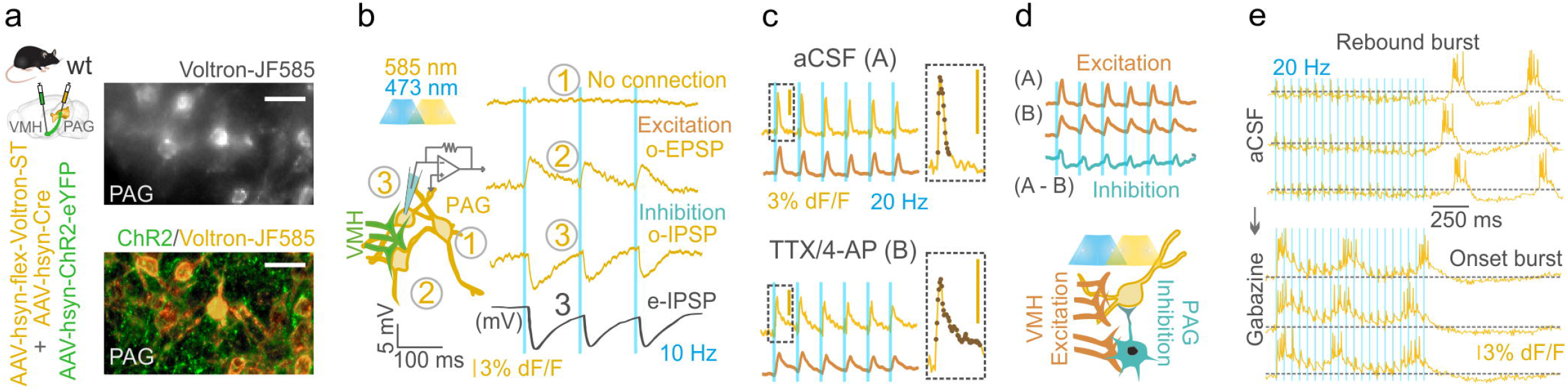
All-optical postsynaptic voltage imaging. All Voltron traces were reversed. **a,** Scheme of viral expression of ChR2 in the VMH-PAG pathway and Voltron-ST in the PAG. Epifluorescent image of neurons with JF-585 signal (top right, scale bar, 30 μm); Confocal image of the same neurons with ChR2 (green) and JF-585-Voltron-ST (gold) labeling in the PAG (bottom right; scale bar, 30 μm) **b,** Scheme of simultaneous all-optical voltage imaging and whole-cell patchclamp recording (left). O-phys traces (average of 7 traces, gold) of three neurons (1,2,3) from top to bottom: neuron with no connection (1) neuron with excitation (o-EPSP) (2) and inhibition (o-IPSP) (3), and e-phys trace (average of 7 traces, black) of the whole-cell-recorded neuron (3) confirmed inhibitory postsynaptic responses (e-IPSP) (right). **c,** All-optical compound o-PSPs in aCSF (A) (top, average of 7 traces, gold) and in TTX (1 μM)/4-AP (5 mM) (B) (bottom, average of 7 traces, gold) with the moving averages below (average of 7 traces, brown). Inserts show the kinetics of the compound signal (top) and o-EPSP (bottom) with the detected datapoints overlayed on the o-phys traces. **d,** Top, subtraction of moving averages of A and B to extract the disynaptic inhibitory signal component (blue) (A-B) eliminated by TTX/4-AP. Scheme of all-optical voltage imaging of a PAG neuron receiving excitation from VMH and disynaptic inhibition from putative local PAG circuitry (bottom). **e,** All-optical voltage imaging of PAG neuron with inhibition during Op of VMH input and rebound burst firing after the Op (top, “Rebound burst”); same PAG neuron in bath-applied γ-aminobutyric acid ionotropic receptor (GABAAR) antagonist, Gabazine (10 μM) showed elimination of the GABA_A_-mediated inhibition resulting in onset burst firing during Op (bottom, “Onset burst”).

### Classification of postsynaptic all-optical response types

To all-optical image the entire VMH-PAG connectome, we designated the PAG area with high density of VMH axons based on a 3D axonal map (Extended Data Fig. 3a). (Fig. 2a). We all-optical imaged 2 to 3 planes in each FOV, each with 30-80 neurons, tile-covered the PAG with 6-7 FOVs (200×350 μm) on 5-7 brain slices per mouse in 7 mice and imaged 6911 VMH-PAG neurons (Fig. 2b, Extended Data Fig. 3c). Somatic region of interests (ROIs) were detected by implementing Cellpose segmentation^13^. Optical-physiology (o-phys) traces were extracted (Fig. 2c) and o-AP peaks, subthreshold (o-Sub) kinetics, o-EPSPs, o-IPSPs and burst activity was detected on each o-phys trace (Fig. 2d). Periods of burst were detected based on o-Sub kinetics and validated by the firing frequency during the detected periods (Extended Data Fig. 3c,d,e). We validated the agreement of bursting PRTs and intrinsic excitability of onset burster PAG neurons (Extended Data Fig. 3f). We designed *Concentric* analysis to validate the somatic origin of o-phys traces based on the o-AP peak amplitudes, and length of bursts (Extended Data Fig. 3g,h,i). Overall, ~89% of the imaged PAG neurons were connected to the VMH based on the detection of >3 o-PSPs, 4% of PAG neurons had o-IPSPs. All-optical testing evoked AP firing in ~67% of PAG neurons (Fig. 2e). To classify the VMH-PAG PRTs, we extracted 29 o-phys parameters (Extended Data Fig. 4a) and performed unbiased hierarchical clustering. We classified the 6911 PRTs into 18 distinct clusters (Fig. 2f,g, Extended Data Fig. 4b,c) and we identified: persistent activity after the Op (cluster 1,4); rhythmic bursting with 3-4 Hz (cluster 8), separate burst upon each Op (cluster 5); time-locked single o-AP upon each Op (cluster 7); strongly depressing (cluster 10,6) and strongly facilitating short-term synaptic plasticity (cluster 9); facilitating paired-pulse (PP) plasticity (cluster 2,4,9,16); depressing PP plasticity (cluster 15); spontaneous firing activity preceding the Op (cluster 3,11,12,13,14); inhibitory responses (cluster 14); inhibitory responses with rebound firing or bursting (cluster 3); subthreshold response (cluster 17); weak or not detectable connection (cluster 18) (Fig. 2h). Suprathreshold PRTs have both pre- and postsynaptic components, report the interaction of these features, and display intrinsic properties (Extended Data Fig. 3f). We validated the lack of animal batch-effect across tSNE regions of clusters (Extended Data Fig. 4d,e). O-phys parameters well-mapped across PRT types which displayed pronounced suprathreshold responses, referred to as “high synaptic drive” (Fig. 2i). For example, spike bimodality was highest in cluster 8, 5; the dynamics in burst length during the Op was increasing in cluster 9 and strongly decreasing in cluster 6; burst length could only be measured in *Post-2* in cluster 1,3,4,11 (Fig. 2i). In summary, our workflow and analysis could capture and resolve the diversity of PRTs with well-defined clusters (Extended Data Fig. f). The VMH-PAG all-optical connectome revealed the total numbers (Extended Data Fig. 4g) and proportions of subthreshold and suprathreshold PRTs across the entire PAG.

**Fig. 2:**
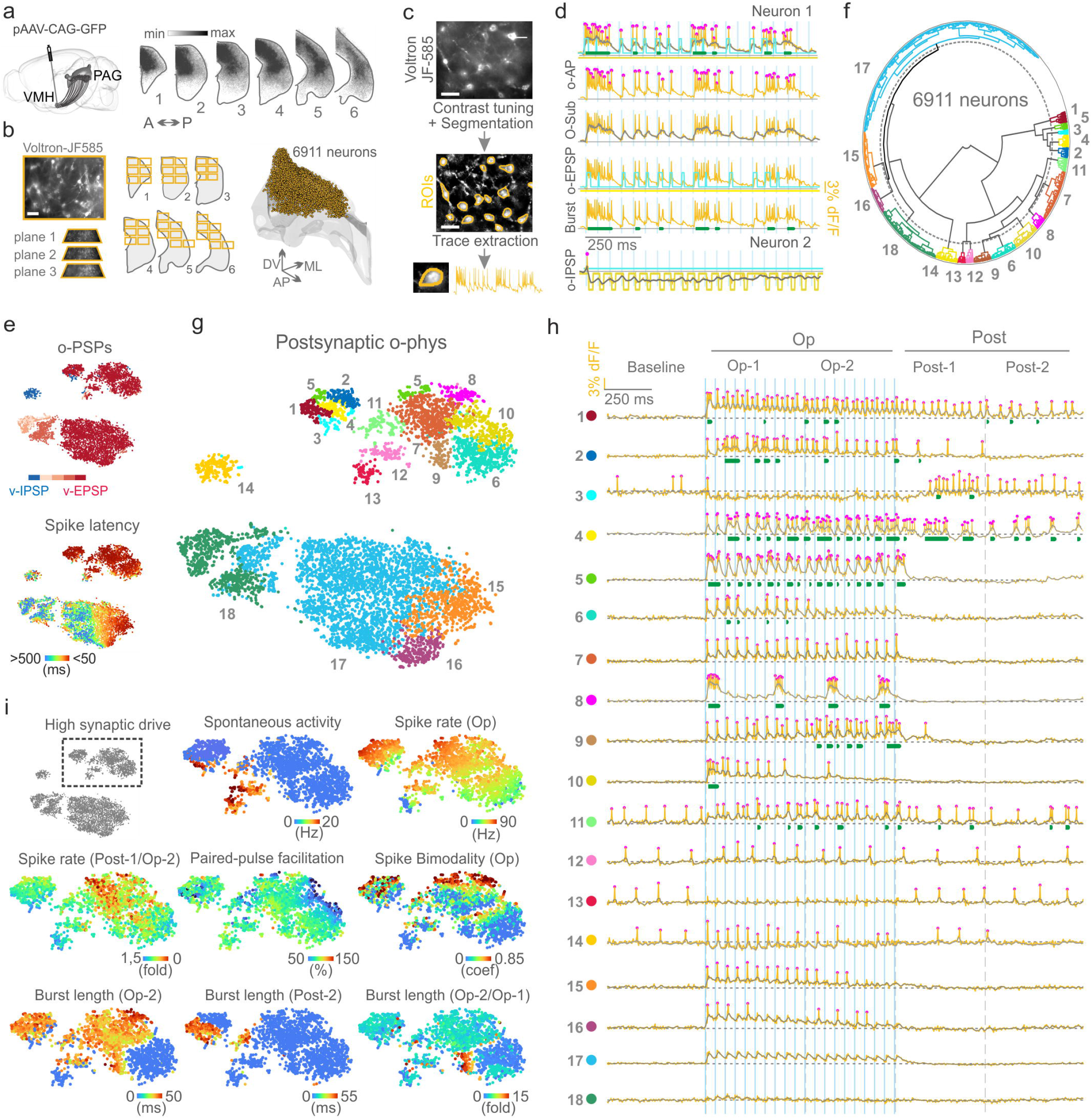
Classification of all-optical postsynaptic response types. All Voltron traces were reversed. **a,** Scheme of viral GFP labeling of VMH-PAG (left) and coronal bins of our 3D axonal map shows PAG areas with high VMH axonal coverage (right). **b,** Example of a FOV with JF-585-Voltron-ST PAG neurons and the illustration of tile-covering PAG slices for all-optical imaging (left) 3D plot of PAG with the 6911 neurons of our all-optical connectome (right). **c,** representative FOV frame average of JF-585-Voltron-ST PAG neurons (top), a contrast-tuned version of the same frame average with the yellow contours of segmented ROIs (middle), example o-phys trace extracted from an ROI (bottom). **d,** Detection of o-APs, O-Sub, Burst activity, o-EPSPs and o-IPSPs. **e,** t-SNE plot of the VMH-PAG o-phys connectome color coded by the average number of detected o-IPSPs (blue) or average number of o-EPSPs (shades of red for ranges: 1-5;5-10;11-15;16-20) (top), same plot color coded by the latency of the first AP detected during the OS (bottom). **f,** Polar dendrogram of agglomerative hierarchical clustering with the identified clusters numbered and colored. **g,** t-SNE plot of the identified o-phys clusters, numbering and color code is identical to f. **h,** Scheme on top details the temporal segments of all-optical sweeps with Op-1: first half of OS, Op-2: second half of OS, Post-1: first half of after-OS, Post-2: second half of after-OS. Representative o-phys traces illustrate PRTs of the identified o-phys clusters (color code and number is identical to f and g) blue bars indicate the 20 Hz 473 nm Op. **i,** t-SNE of the VMH-PAG postsynaptic o-phys (gray), crop of “High synaptic drive” clusters, which are enlarged in the following t-SNEs mapping clustering parameters. Abbreviation in brackets for each parameter indicates the temporal segments of the parameter extraction.

### Spatial topography of postsynaptic connectome

We post-hoc extracted the XYZ coordinates of each 6911 imaged neuron and spatially mapped the VMH-PAG connectome (Extended Data Fig. 5a,b). For spatial PRT-independent overview, the PAG was divided to voxels, and in each, the percentage of PRTs fulfilling a criterion was calculated (Fig. 3a). The voxel-criterion of PRTs with >3 o-PSPs displayed a homogenous distribution of high-percentage voxels confirming complete coverage of connections throughout the VMH-PAG connectome (Fig. 3b). The voxel-criterion of PRTs with >18 o-PSPs visualized stronger connections in a spatial pattern (Fig. 3c) capturing the more anterior and lateral PAG volumes with higher VMH synaptic drive. Voxel-mapping paired-pulse facilitation, a qualitative short-term synaptic plasticity property^14^, highlighted PAG subregions with high density of facilitating connections (Fig. 3d). Remarkably, mapping the VMH-PAG connectome by o-phys cluster identity revealed spatial topography of PRT clusters (Fig. 3e). The coverage of d-dl-lPAG allowed to inspect the distribution of clusters across these areas (Extended Data Fig. 5c,d). To test the spatial proximity within and across o-phys clusters, the average minimal distance (AMD) between neurons of the same cluster was compared to the chance-level-AMD calculated by repeated shuffles of cluster identity (Fig. 3f). The observed AMDs were shorter than chance-level for most clusters, thus o-phys clusters also formed spatial clusters. The AMD of neurons across the 18 o-phys clusters was significantly smaller than chance-level between spatially intermingled clusters 2 and 4, cluster 10 and 1, and cluster 9 and 6 (Fig. 3g). The AMD was significantly larger than chance-level between spatially segregated clusters, such as cluster 9 and 13 or cluster 8 and 1 (Fig. 3g). The side-by side visualization of o-phys cluster centroid distances and spatial cluster centroid distances suggested coherence of the two properties (Fig. 3h). The function of the two distances confirmed a pronounced correlation between o-phys cluster identity and spatial location (Fig. 3i). Taken together, our all-optical connectome described whole-structure VMH-PAG connectivity on the quantitative and qualitative levels. Spatial mapping of our whole-structure VMH-PAG connectome could reveal the topography of PRT clusters using 7 mice. This throughput was not accessible by any other approach to-date.

**Fig. 3:**
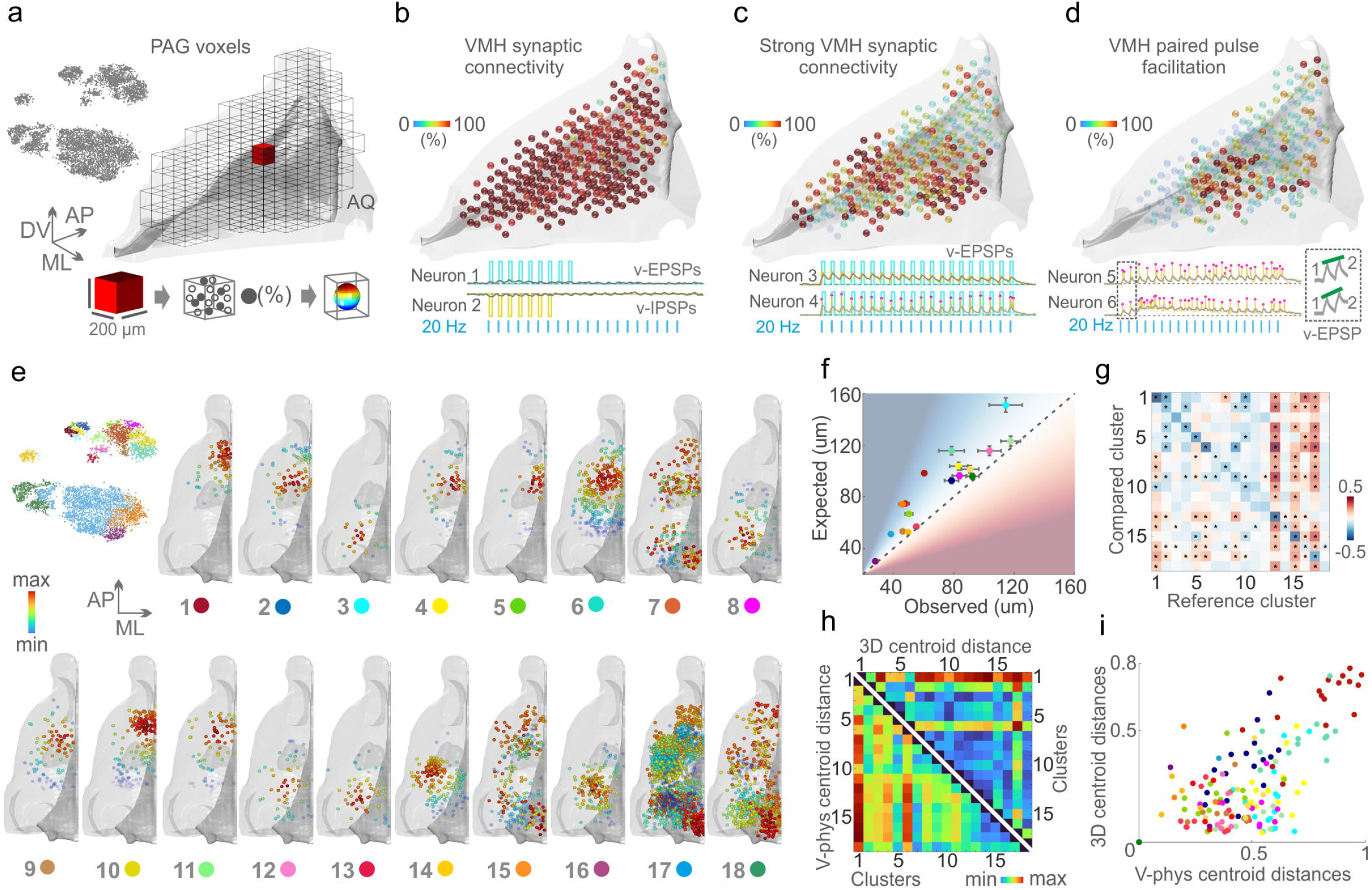
Spatial topography of postsynaptic connectome. **a,** t-SNE of the VMH-PAG connectome in gray, no cluster information was used (top left). 3D PAG model of hemisphere subdivided to voxels (grid) with one voxel in the middle (red). Illustration of a 200×200 μm voxel and the neurons inside the voxel fulfilling a criterion that defined the color code from 0% (blue) to 100% (red) and the transparency with 0% being invisible (bottom). **b,** 3D PAG model with voxel-mapping of overall VMH connectivity (top), example PRTs fulfilling the low-cut criterion of displaying >3 PSPs (Neuron1,2). **c,** 3D PAG model with voxel-mapping of strong VMH synaptic connectivity (top), example PRTs fulfilling the low-cut criterion of displaying >18 PSPs (Neuron3,4). **d,** 3D PAG model with voxel-mapping paired-pulse facilitation (PP) (top), example PRTs of postsynaptic neurons exerting facilitating PP, insert shows first two o-EPSPs with green line highlighting the facilitation (Neuron5,6). **e,** t-SNE of the VMH-PAG connectome color coded by cluster identity (Fig. 2g,h) (top left). 3D PAG model of one hemisphere for each o-phys cluster with the spatial density-core mapping. Colors from red to blue go from highest to lowest spatial density, transparency follows the color code with blue being invisible. **f,** observed average minimal distance (AMD) between neurons of the same cluster, versus the AMD expected by chance calculated by repeated shuffles of cluster identity (error bars: ±sem (bootstrapped)). **g,** AMD of neurons of distinct o-phys clusters was probed across all 18 clusters (* = p-value < 3*10^-5^). **h,** side-by side visualization of o-phys cluster centroid distances and spatial cluster centroid distances, all axes are cluster labels. **i,** Scatter plot of o-phys cluster centroid distances versus spatial cluster centroid distances, colors code cluster identity.

### On-site analysis of all-optical voltage imaging with VoltView

To make all-optical experimenting interactive, we developed VoltView. The *Detailed* configuration runs thorough analysis to validate signal source with Concentric analysis, extracts o-AP peaks, O-Sub, periods with burst activity (Burst), o-AP half-width and detects o-EPSPs and o-IPSPs in a FOV in ~4-5 min (Fig. 4a). The *On-site* configuration of the package provides quick access to the basic analyzed features (o-APs, O-Sub, Burst) of 30-80 neurons in ~1-1.5 min (Fig. 4a). After onsite analysis, VoltView opens a “ROI Explorer” for browsing PRTs. The position of the currently inspected ROI is indicated in the FOV to spatially guide the experimenter for further investigation or soma harvesting (Fig. 4b). On-site analysis can compare analyzed features across different all-optical recordings of the same neurons. Using VoltView, we attempted to identify long-term changes of PRTs with increased or decreased burst plasticity evoked by a 50 Hz optogenetic high-frequency stimulation (oHFS) (Fig. 4c). After motion correction of the re-imaged FOVs, parameters of the same neurons were extracted and on-site-compared across imaging sessions (Rec1, Rec2). On-site analysis had a 25%-change cutoff to identify ROIs with increased or decreased total burst length (Fig. 4c). VoltView successfully identified a neuron with robustly decreased bursting where slow bursting turned into continuous firing after oHFS (Fig. 4d,f). A neighboring neuron displayed opposite change with increased burst length where interestingly the rhythmicity of bursts blended into a tone, well-shown by the sweep average (Fig. 4e,f). Importantly, the responses of both example neurons in both Rec1 and Rec2 were highly stable across 7 sweeps taken during a 70 s time window, indicating that all-optical observation of neurons can capture a PRT rendered to a circuit state. Altogether, VoltView analysis made *ex vivo* all-optical imaging interactive by on-site selection of neurons with specific PRT based on user-defined firing property or connection plasticity features.

**Fig. 4:**
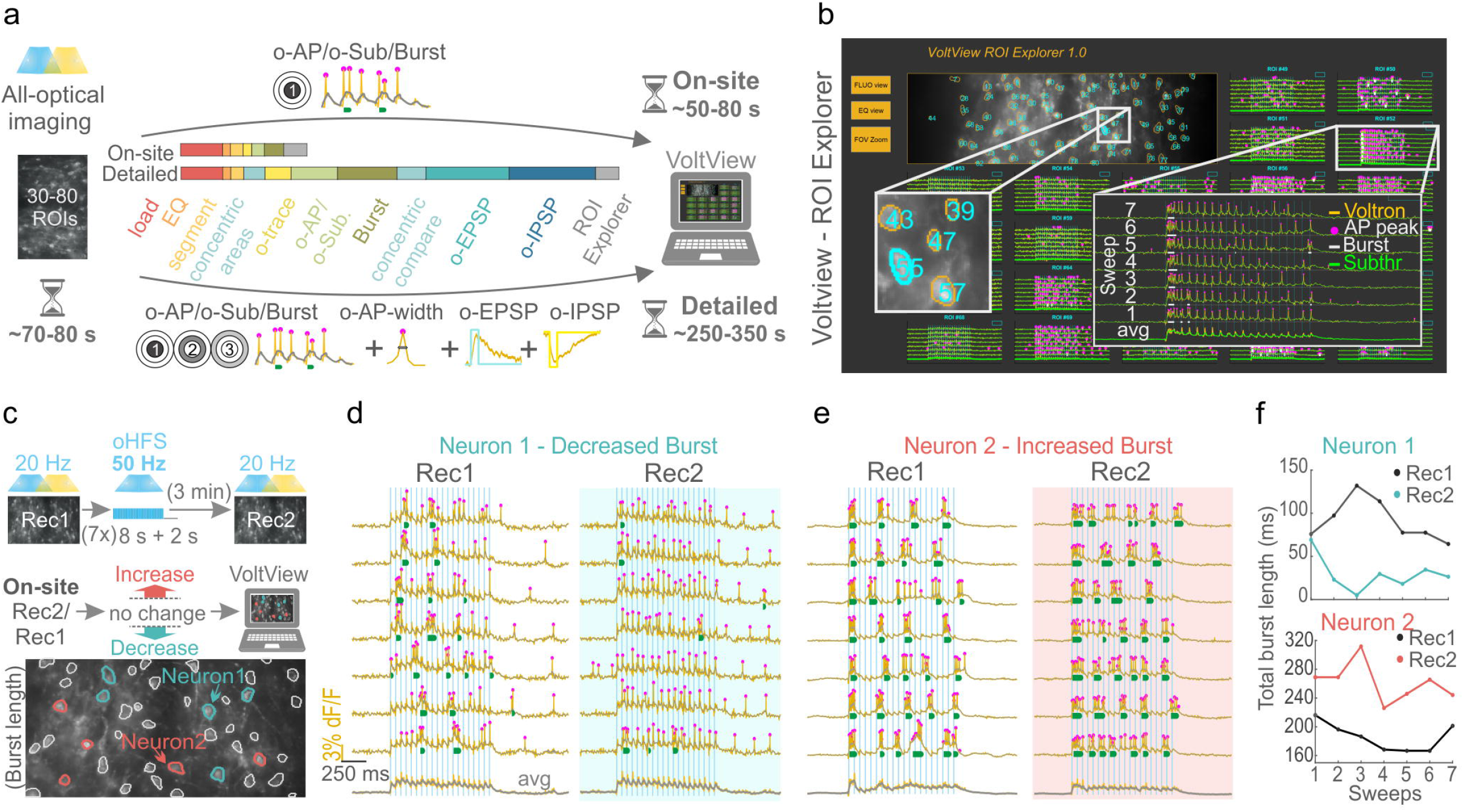
On-site analysis of all-optical voltage imaging with VoltView. All Voltron traces were reversed. **a,** Illustration of the *On-site* (top arrow) and *Detailed* (bottom arrow) analysis of o-phys. Logos summarize the analyzed features on the arrows. Colored bars (red to blue) compare the proportion of time for different processing, detection and analysis modules **b,** VoltView with the ROI explorer. Enlarged insert demonstrates ROI spatial indication with the number of the corresponding all-optical recording (left). Enlarged insert shows the recorded o-phys sweeps with the detected spikes and bursts (right). **c,** Rec1 and Rec2 repeated recordings in the same FOV, before and after 50 Hz oHFS; VoltView extracts parameters of the same neurons and compares them across Rec1 and Rec2. Decreased bursting is indicated with blue ROI contours, increased bursting is indicated with red ROI contours (bottom). **d,** Neuron1: neuron which initially responded with bursts to the 20 Hz Op and after 50 Hz oHFS the response shifted towards continuous firing (blue). Bottom sweep is the average of the 7 sweeps above. **e,** Neuron2: initially exerted rhythmic bursts during Op and after the 50 Hz oHFS responded to the 20 Hz Op with prolonged bursts, losing rhythmicity. The last sweep is the average of the 7 sweeps above. **f,** Line plots show comparison of total burst lengths before (black) and after oHFS (Neuron1 blue, Neuron2 red). Dots represent the total burst lengths in every sweep.

### VoltView-guided Voltage-Seq

High-throughput all-optical connectivity testing probes dozens of connections simultaneously, up to 1000-1500 the same day (Fig. 5a). To on-site-navigate to specific PRTs in such a large data stream, we added a classifier to VoltView with our VMH-PAG connectome data as a reference (Fig. 5b). VoltView suggested neurons classified into user-defined PRT clusters (Fig. 5c) for soma harvesting (Fig. 5d) and for subsequent scRNA-seq (Fig. 5e). We named the workflow *Voltage-Seq*. To test Voltage-Seq, we attempted to find sparse GABAergic neurons within the VMH-PAG connectome. GABAergic neurons of d-dlPAG had been reported to be depolarized and often spontaneously firing to provide tonic inhibition^15^. Cluster 3,13 and 12 had 42%,95% and 100% spontaneous firing (Extended Data Fig. 6a), were denser in the posterior PAG in our data (Extended Data Fig. 5b), and localized in the d-dlPAG, similarly to the distribution of PAG GABAergic neurons (Extended Data Fig. 6c,i).To probe the correlation of GABAergic identity and PRTs of cluster 3,12 and 13, we all-optical imaged 1436 GABAergic neurons in the posterior PAG of 4 VGAT-Cre mice. Cluster-load analysis of VGAT data confirmed large wild-type cluster-load in cluster 3,12, and 13 (Extended Data Fig. 6a,d,e). We set VoltView to suggest neurons of cluster 3,12 and 13 during all-optical imaging of VMH-PAG PRTs and harvested 60 neurons from 3 wildtype mice. The harvesting protocol was optimized for Voltage-Seq (see *Methods*), for scRNA-seq we used Smart-Seq2^16^. Noise-level detection of glial markers and the detection of ~6000 genes/cell confirmed high RNA-transcriptome quality (Extended Data Fig. 6f,g,h). The chance-level of finding GABAergic vs. Glutamatergic neurons was estimated to be ~22% based on in situ hybridization (ISH) labeling density of GABAergic (*Slc32a1*, *Gad1*, *Gad2*) vs. glutamatergic (*Slc17a6*) molecular markers (Fig. 5g, Extended Data Fig. 6i,j). Unbiased clustering of Voltage-seq (Fig. 5f) showed 3-times higher, 72% (39/54) ratio of GABAergic neurons (Fig. 5g,h), significantly more than expected to find by chance. Remarkably, amongst the GABAergic PRTs we found a circuit phenomenon which had not yet been described before. We observed PRTs we named *Switch* response, where converging excitatory and inhibitory inputs dominated in a switching manner. The Switch behavior was stable and reproducible (Extended Data Fig. 7a). Gabazine eliminated the inhibitory input and turned every response excitatory (Fig. 5i). Taken together, Voltage-Seq successfully identified sparse GABAergic neurons of the VMH-PAG connectome based on the on-site classification in VoltView and could provide access to molecular identity of the same neurons. Furthermore, during this test, our methodology revealed a hitherto unknown disinhibitory circuit motif.

**Fig. 5:**
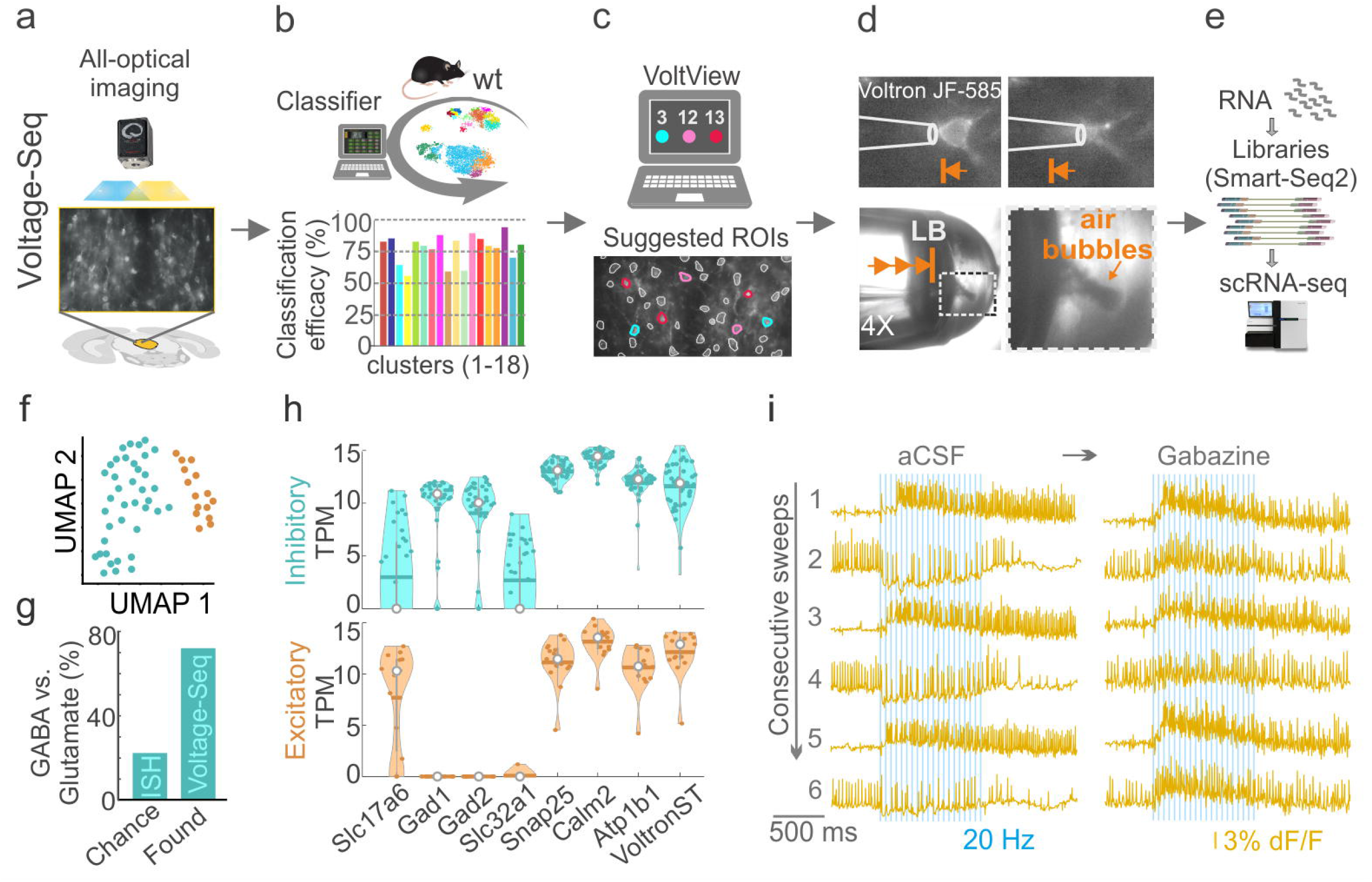
VoltView-guided Voltage-Seq. **a,** Voltage-Seq workflow: all-optical voltage imaging; **b,** on-site analysis with classification (classifier efficacy for each cluster on the bar plot color coded by cluster identity); **c,** VoltView suggests ROIs; **d,** soma harvesting (upper row) in 20X magnification with ejection (bottom row) under 4X magnification validated by air bubbles upon ejection of sample in the lysis buffer (LB); **e,** cDNA library preparation and scRNA-seq. **f**, UMAP of unbiased clustering of Voltage-Seq RNA-transcriptome (inhibitory neurons: blue, excitatory neurons: brown). **g**, Bar plot comparing the proportion of GABAergic versus glutamatergic neurons in our sample (Voltage-Seq, Found) compared to the chance-level (ISH, Chance). **h**, Violin plots show log2 TPM levels of neurotransmitter modality defining *Slc17a6*(Vglut2), *Gad1* (GAD 67), *Gad2* (GAD 65), *Slc32a1*(Vgat) and pan neuronal control genes *Snap25* (Synaptosomal-Associated Protein 25), *Calm2* (Calmodulin 2), *Atp1b1* (Sodium/potassium-transporting ATPase subunit beta 1), *Voltron-ST* (somatargeting Voltron) in the GABAergic (Inhibitory, blue) and glutamatergic (Excitatory, brown) clusters; horizontal bars represent mean, white dots represent median. **i**, consecutive sweeps of a Switch response in aCSF (left) and in Gabazine (right)where all sweeps turned excitatory.

### Neuronal identity and neuromodulation in the Switch motif

Voltage-seq unveiled that a Switch motif regulates the activity of certain GABAergic (4/39) neurons in the d-dlPAG (Extended Data Fig. 8a). To examine the postsynaptic neuronal identity in the newly found Switch motif, we all-optical imaged VMH-PAG in wild-type mouse and browsed the PRTs in VoltView to find Switch responses (Fig. 6a). After locating a Switch response (Fig. 6b), we whole-cell-recorded the postsynaptic neurons. We characterized the intrinsic excitability and remarkably found both burst firing (6/8) and regular spiking (2/8) types (Fig. 6c). Thus, Switch motif is not rendered to one specific neuronal type but is a circuit domain integrating multiple neuronal types. To confirm the disynaptic nature of inhibition we measured 4.86±2.36 ms IPSC delay in burster and 4.72±1.78 ms IPSC delay in regular spiking neurons in voltage-clamp mode during Op (Fig. 6d). Furthermore, we probed GABAergic identity with *Slc32a1* ISH and found that all the investigated neurons with Switch response were GABAergic, in agreement with our Voltage-seq transcriptome data (Fig. 6e). Differential expression (DE) analysis of excitatory and inhibitory clusters highlighted putative marker genes of GABAergic neurons (e.g. *Nrxn3*, *Pnoc*, *Gata3*) (Fig. 6f, Extended Data Fig. 8c). Interestingly, based on our Voltage-seq RNA-transcriptome, amongst other GABAergic cells, the neurons with Switch response were expressing *Tacr1* and *Tacr3* encoding neurokinin-1 and −3 (NK-1 and NK-3) receptors (Fig. 6f, Extended Data Fig. 8b). To probe neuromodulation of the Switch motif mediated by NK-1 and NK-3, we bath-applied Substance P (SP), the endogenous ligand of the receptors^17^. Multiple video comparisons in VoltView showed increased PRT firing frequency only in a subset of neurons (5/58) (Fig. 6i). In a strong Switch responder with a rebound burst only after excitatory sweeps, we could voltage image SP neuromodulation (Fig. 6g,h). Subthreshold excitatory responses turned suprathreshold, while inhibitory responses were suppressed and rebound bursts were eliminated. This could be due to the SP-induced inward Na^+^-current increasing the firing probability and counteracting inhibition^18^. Furthermore, we located this neuron in VoltView, and with whole-cell recording we identified a burst firing type, which could exert rebound burst firing (Fig. 6j). We successfully validated the GABAergic identity of the same neuron with *Slc32a1* ISH (Fig. 6k). In summary, our Voltage-seq methodology could allow closer investigation of a newly described disynaptically disinhibitory circuit motif. Switch responders could not have had been found by whole-cell patch-clamp as they are relatively sparse. Based on the Voltage-seq transcriptomics we could identify SP as a neuromodulator of the Switch motif, and all-optical image the SP-neuromodulation of the Switch motif.

**Fig. 6:**
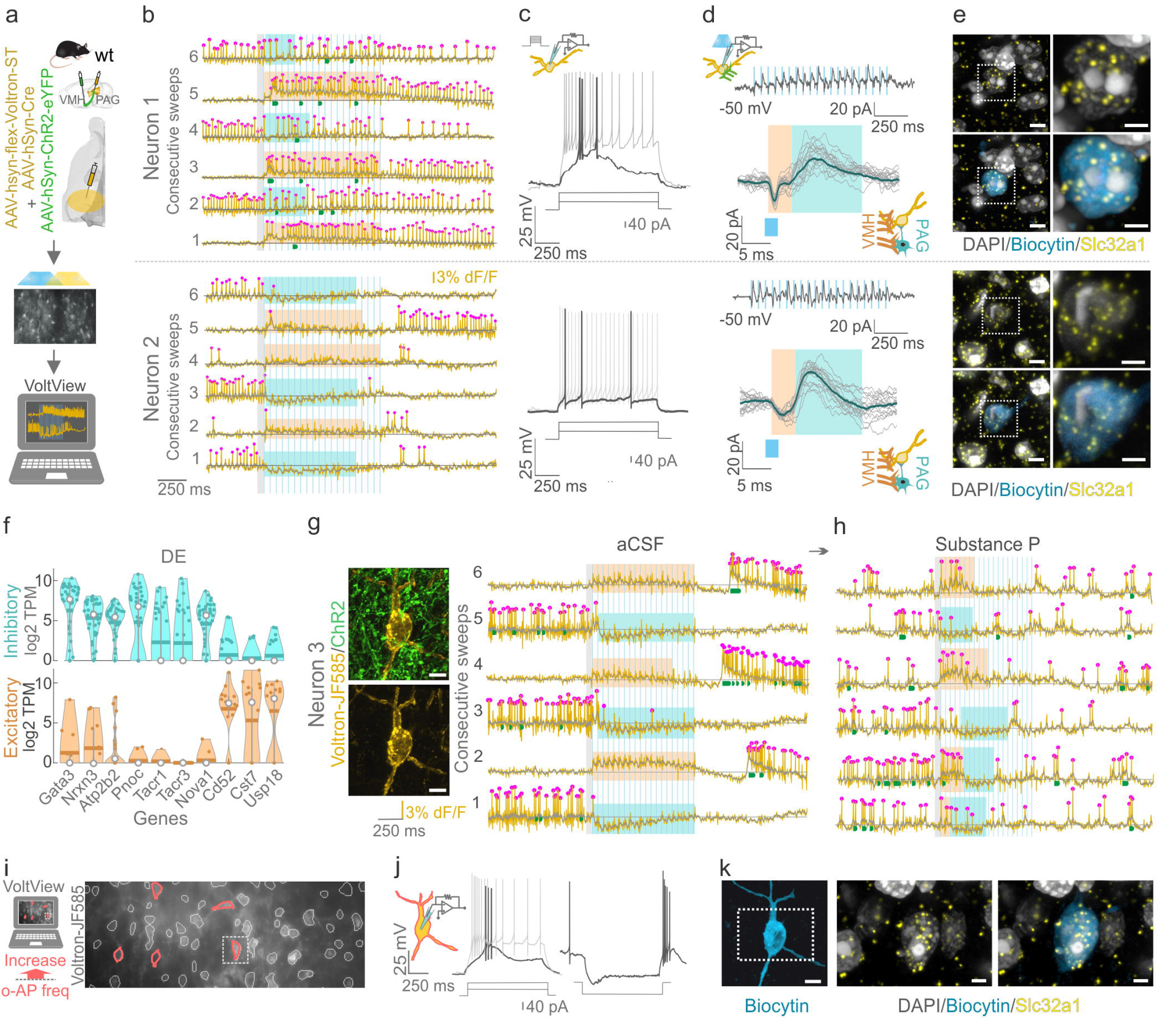
Neuronal identity and neuromodulation in the Switch motif. All Voltron traces were reversed. **a,** Scheme of viral expression of ChR2 in the VMH-PAG pathway and Voltron in the posterior PAG (top). All-optical voltage imaging (middle) and on-site analysis in VoltView (bottom). **b,** O-phys traces of two neurons Neuron1 (top), Neuron2 (bottom) with Switch response. Rectangles represent the overall inhibition (below baseline, cyan) and excitation (above baseline, brown). **c,** Whole-cell current-clamp recording shows the bursting firing type of Neuron 1 (top) and regular spiking type of Neuron 2 (bottom). **d,** Neuron 1(top panel): voltage-clamp trace of current responses upon 20 Hz Op of VMH terminals (−50 mV, average of 7 sweeps) (top). Overlayed interstimulus intervals with e-EPSCs (brown rectangle) and e-IPSCs (cyan rectangle) demonstrate the delay of evoked responses during Op (below); 1.52±0.2 ms EPSC delay in bursters, 1.86±0.73 ms EPSC delay in regular spikers. Same experiment for Neuron 2 (bottom panel). **e,** ISH validation of Slc32a1 positivity of Neuron1 (top panel). DAPI (white) and Slc32a1 (yellow) colocalization (upper row). DAPI (white), Slc32a1(yellow) and Biocytin (blue) colocalization (lower row) (scale bar, 5 μm, inserts 2 μm). Insert shows Neuron1 in higher magnification. Same experiment for Neuron2 (bottom panel). **f,** Violin plot of differentially expressed genes of GABAergic (cyan) and glutamatergic (brown) neurons. *GATA3* (GATA Binding Protein 3), *Nrxn3* (Neurexin 3), *Atp2b2* (ATPase Plasma Membrane Ca^2+^ Transporting 2), *Pnoc* (Prenociceptin), *Tacr1* (Tachykinin Receptor 1),*Tacr3* (Tachykinin Receptor 3), *Nova1* (NOVA Alternative Splicing Regulator 1), CD52 (Campath-1 antigen), *Cst7* (Cystatin F),*Usp18* (Ubiquitin specific peptidase 18). **g,** JF585-Voltron labeled (gold) PAG neuron (Neuron3) surrounded by ChR2-expressing axons (green) (left) (scale bar, 6 μm). Consecutive o-phys traces from the same neuron with a strong Switch response (aCSF). Rectangles represent the overall inhibition (below baseline, cyan) and excitation (above baseline, brown). **h,** Consecutive o-phys traces of the same neuron after bath-application of SP (1 μM). Rectangles represent the overall inhibition (below baseline, cyan) and excitation (above baseline, brown). **i,** Multiple video comparison in VoltView indicated neurons of the FOV that responded with increased AP firing upon SP wash-on (red ROIs). Insert (dashed line) marks the recorded Switch responder from g,h. **j,** Scheme of whole-cell recording of the Switch responder Neuron3 modulated by SP (left). Whole-cell current clamp recording revealed the bursting firing type of the Switch responder (middle), which exerted rebound burst firing after negative current injection (right). **k,** whole-cell recorded biocytin-filled Neuron3 from g,h,i,j (blue, left). Inserts show the DAPI (white) and *Slc32a1* (yellow) colocalization (left) and the DAPI (white), *Slc32a1* (yellow) and biocytin (blue) colocalization (right) in the same Switch responder Neuron3.

## Discussion

We optimized Voltage-Seq which combines, all-optical physiology, spatial mapping, on-site classification, and RNA-transcriptomics to robustly increase the throughput of synaptic connectivity testing and targeted molecular classification of postsynaptic neurons. The literature of VMH-PAG synaptic physiology is yet exiguous. Our approach provided a detailed insight into the VMH-PAG synaptic connectome with measuring thousands of connections on the qualitative and quantitative levels using only a few animals. Such connectome data in any brain region is a highly potent starting point for addressing questions concerning specific PRTs and the postsynaptic neuronal types.

Comprehensive understanding of the identity of postsynaptic ensembles was so far occluded by the throughput of connectivity probing techniques. Besides, patch-clamp may introduce excess perturbation of the intracellular ionic milieu, for example we could not induce the switching behavior of a Switch responder during whole-cell recording. In contrast, Voltage-Seq of circuit motifs could capture the native circuit behavior of the integrated neurons. The interplay of long-range synaptic inputs, local connectivity, and intrinsic properties of postsynaptic neurons could be observed and Voltage-Seq could resolve the molecular identity of these neurons.

With the help of such a multilayered methodology, researchers could address more complex functional connectivity questions. For example, high-throughput connectome analysis in mouse models of mood disorders could provide an overview of the short-term and long-term synaptic plasticity landscape of one or multiple brain areas of the same mouse. Such a dataset could highlight the PRTs of neurons in the focal points of circuit imbalance. Voltage-Seq of these target neurons could reveal the identity and detect gene expression changes of postsynaptic neurons in circuit motifs of imbalanced networks.

Voltage-Seq methodology will be more accessible to a wider range of neuroscientists as it needs less hands-on skills compared to patch-clamp-electrophysiology which requires years of dedicated practice. Voltage-Seq united the power of high-throughput, high-signal-resolution connectivity imaging, interactive analysis and molecular profiling of the imaged neurons which is a major advancement in postsynaptic circuit dissection.

## Methods & Protocols

### Animals

Experiments were conducted using adult male and female mice, wild type C57BL/6J (Charles River Laboratories) or the transgenic mouse line VGAT-Cre: *B6J.129S6(FVB)-Slc32a1^tm2(cre)Lowl^/MwarJ*, Jackson stock no. 028862. All transgenic mice used in experiments were heterozygous for the transgenes. Mice were group housed, up to five per cage, in a temperature (23°C) and humidity (55%) controlled environment in standard cages on a 12:12hr light/dark cycle with *ad libitum* access to food and water. All procedures were approved and performed in accordance and compliance with the guidelines of the Stockholm Municipal Committee (approval no. N166/15 and 7362-2019).

#### Animal cohorts

Axonal anatomy and histology: n = 2 pAAV-CAG-GFP

Patch-clamp electrophysiology: n = 7 pAAV-hSyn-hChR2(H134R)-EYFP

Voltage-imaging and optimalization: n = 21 pENN-AAV-hSyn-Cre-WPRE-hGH; pAAV-hsyn-flex-Voltron-ST; pAAV-hSyn-hChR2(H134R)-EYFP

### Viral constructs

All purified and concentrated adeno-associated viruses (AAV) were purchased from Addgene.

#### Anatomy and histology

pAAV-CAG-GFP (AAV5); Addgene, cat. no. 37825-AAV5; (at titer ≥ 7×10e12 viral genomes/ml)

#### Voltage-imaging

pAAV-hsyn-flex-Voltron-ST (AAV1); Addgene, cat. no. 119036-AAV1; (at titer ≥ 2×10e12 viral genomes/ml)

pAAV-hSyn-hChR2(H134R)-EYFP (AAV5); Addgene, cat. no. 26973-AAV5; (at titer ≥ 7×10e12 viral genomes/ml)

pENN-AAV-hSyn-Cre-WPRE-hGH (AAV1); Addgene, cat. no. 05553-AAV1; (at titer ≥ 1×10e13 viral genomes/ml)

### Viral injections

#### General procedure

Mice were anesthetized with isoflurane (2%) and placed into a stereotaxic frame (Harvard Apparatus, Holliston, MA). Before the first incision the analgesic Buprenorphine (0.1 mg/kg) and local analgesic Xylocain/Lidocain (4 mg/kg) was administered subcutaneously. The body temperature of the mice was maintained at 36 °C with a feedback-controlled heating pad. For viral injections a micropipette attached on a Quintessential Stereotaxic Injector (Stoelting, Wood Dale, IL) was used. Injections were done with the speed of 50 nl/minute. The injection pipette was held in place for 5 min after the injection before being slowly (100 μm/s) retracted from the brain. The analgesics Carprofen (5 mg/kg) was given at the end of the surgery, followed by a second dose 18-24h after the surgery.

#### Labeling strategies

For anatomical characterization and electrophysiological recordings of the VMH-PAG pathway in C57BL/6J mice, 0.3μl pAAV-CAG-GFP or pAAV-hSyn-hChR2(H134R)-EYFP was unilaterally injected into the VMH (coordinates: AP −1.45 mm, ML 0.25 mm, DV −5.25 mm). Targeting of the PAG was achieved by 1 (coordinates: AP −4.1 mm, ML 0.2 mm, V −1.6 mm) or 2 (coordinates: AP −3.9 mm and AP −4.3 mm, ML 0.2 mm, V −1.6 mm) unilateral injections of 0.3μl of 1:1 mixture of pENN-AAV-hSyn-Cre-WPRE-hGH and pAAV-hsyn-flex-Voltron-ST.

### Histology

#### General procedure

Mice were deeply anaesthetized with Na-pentobarbital (60mg/kg) and transcardially perfused with 0.1M PBS followed by 4% paraformaldehyde in PBS 0.1M. Brains were removed and post-fixed in 4% paraformaldehyde in PBS 0.1M overnight at 4°C and then washed and stored in 0.1M PBS. Coronal, 50 μm slices were cut using a vibratome (Leica VT1000, Leica Microsystems, Nussloch GmbH, Germany). The sections were washed in 0.1M PB and mounted on glass slides (Superfrost Plus, Thermo Scientific) and coverslip-covered(Thermo Scientific) using glycerol: 1x PBS (50:50).

#### Histology of biocytin filled neurons

250 μm thick brain slices containing biocytin-filled neurons and voltron-JF585 labeling were postfixed in 4% paraformaldehyde in phosphate-buffer (PB, 0.1M, pH 7.8) at 4 °C overnight. Slices were repeatedly washed in PB and cleared using CUBIC protocol^19^. First “CUBIC reagent 1” was used (25 wt% urea, 25 wt% N,N,N’,N’-tetrakis(2-hydroxypropyl) ethylenediamine and 15 wt% polyethylene glycol mono-p-isooctylphenyl ether/Triton X-100) for 1 day at 4 °C. After repeated washes in PB, biocytin was visualized using Alexa Fluor 633-conjugated streptavidin (1:1000, RT 3 hours). For NeuN staining primary antibody (Millipore, MAB377,1:1000) was incubated overnight at 4 °C and after repeated washing with PB, 2nd antibody (Jackson, Cy5, Code:715-175-151) was incubated for 3h RT. Slices were then re-washed in PB and submerged in “CUBIC reagent 2” (50 wt% sucrose, 25 wt% urea, 10 wt% 2,20,20’-nitrilotriethanol and 0.1% v/v% Triton X-100) for further clearing. Slices were mounted on Superfrost glass (Thermo Scientific) using CUBIC2 solution and covered with 1.5 mm cover glasses.

#### In situ hybridization (ISH)

We used *RNAscope Fluorescent Multiplex Assay V2* (cat.no. 323110) to visualize *Slc32a1* in biocytin-filled, voltage-imaged neurons. Brain slices after all-optical voltage imaging were fixed overnight at 4°C in 4% paraformaldehyde, next day slices were repeatedly washed in PB. FISH protocol followed the manufacturer instructions with modified incubation time as slices were 250 μm thick. We incubated the free-floating slices with the ISH probe overnight instead of 2 h on a slide, at 40°C, next day washed the sections in wash buffer and used AMP-1FL for 30 min at 40°C and Amp-2FL for 15 min at 40°C. Jf-585 labeling/fluorescence got almost eliminated by the ISH protocol, thus we also took images before and after ISH. We attempted to re-incubate PFA-fixed tissue in JF-585-HaloTag-dye but it did not re-label the Voltron-ST-expressing neurons. Immunostaining of biocytin worked both before and after ISH, for this we used the same protocol as above (*Histology of biocytin filled neurons*) without the CUBIC clearing steps.

#### Confocal imaging

All confocal images were taken using a Zeiss 880 confocal microscope. CUBIC cleared sections after slice electrophysiology and biocytin or NeuN staining were acquired as z-stacks using a Plan-Apochromat 20x/0.8 M27 objective (imaging settings: frame size 1024×1024, pinhole 1AU, Bit depth 16-bit, speed 6, averaging 4). For viral expression overview of coronal cut VMH or PAG sections were acquired with a Plan-Apochromat 20x/0.8 M27 objective (imaging settings: frame size 1024×1024, pinhole 1AU, Bit depth 16-bit, speed 7, averaging 2). For ISH images oil-immersion 63x/1.0 objective was used (imaging settings: frame size 1024×1024, pinhole 1AU, Bit depth 16-bit, speed 6, averaging 4) Processing of images was either done in ImageJ (NIH, USA) or Imaris 7.4.2 (Oxford Instruments, UK).

### Brain slice preparation *ex vivo*

First, mice were anesthetized with intraperitoneal injection of 50 μl Na-pentobarbital (60 mg/kg) and transcardially perfused with 4-8 °C cutting solution, containing (in mM): 40 NaCl, 2.5 KCl, 1.25 NaH2PO4, 26 NaHCO3, 20 glucose, 37.5 sucrose, 20 HEPES, 46.5 NMDG, 46.5 HCl, 1 L-ascorbic acid, 0.5 CaCl2, 5 MgCl2. Next, brain was carefully removed and 250 μm thick coronal slices were cut with a vibratome (VT1200S, Leica, Germany) in the same 4-8 °C cutting solution. Next, slices were incubated in cutting solution at 34°C for 13 minutes, and kept until recording at room temperature in artificial cerebrospinal fluid (aCSF) solution containing (in mM): 124 NaCl, 2.5 KCl, 1.25 NaH2PO4, 26 NaHCO3, 20 glucose, 2 CaCl2, 1 MgCl2. For Voltron-imaging, slices were incubated at room temperature in Janelia-Fluor 585 HaloTag (JF-dyes, Janelia) ligands. JF-dyes were dissolved in DMSO to a stock of 1 μM and further diluted to 50 nM in aCSF before use. All solutions were oxigenated with carbogen (95% O2, 5% CO2). All constituents were from Sigma-Aldrich.

### Patch-clamp electrophysiology

C57BL/6J mice were injected with pAAV-hSyn-hChR2(H134R)-EYFP at 10-11 weeks and recorded 12-13 weeks of age (for details on the virus injection see “Viral injections” above. For patch-clamp recordings, brain slices were superfused with 33-34°C aCSF at a rate of 4-6 ml/min. Neurons were visualized using a 60x water-immersed objective (Olympus, Tokyo, Japan) in a DIC differential interference contrast (DIC) microscopy on an Olympus BX51WI microscope (Olympus, Tokyo, Japan). Patch borosilicate (Hilgenberg, Germany) pipettes, 7-10 MΩ pulled using a horizontal puller (P-87 Sutter Instruments, Novato, CA, USA) were filled with K-gluconate-internal solution containing (in mM): 130mM K-gluconate, 5mM KCl, 10mM HEPES, 10mM Na2-phosphocreatine, 4mM ATP-Mg, 0.3mM GTP-Na, 8 biocytin, 0.5 ethyleneglycol-bis(2-aminoethylether)-*N*,*N*,*N*’,*N*’-tetraacetate (EGTA), (pH 7.2 set with KOH). The same intracellular solution was used for both current-clamp and voltage-clamp recordings. Signals were recorded with an Axon MultiClamp 700B amplifier and digitized at 20 kHz with an Axon Digidata 1550B digitizer (Molecular Devices, San Jose, CA, USA). Pipette capacitance was compensated, liquid junction potential was not corrected. Neurons recorded in current-clamp mode were held at a membrane potential of −60 mV. “AP threshold” (mV) was defined as the voltage point where the upstroke’s slope trajectory first reached 10 mV/ms. “AP half-width” (ms) was measured at half the maximal amplitude of the AP. To assess the firing types, were held the neurons at a membrane potential of −70 mV and 1 second long positive current was injected. To test the ability of neurons to rebound burst, neurons were held at −60 mV and 1 s long negativ current step was injected. The synaptic properties of VMH projection onto PAG neurons were tested in both voltage- and current-clamp mode on different holding potentials (−70, −60, −50 mV) using multiple light pulse train protocols with 2 ms blue light pulses with ~2.5 mW light power from a SpectraX (Lumencor, USA) LED light source. In some experiments we bath applied TTX (1μM; Tocris) and 4-AP (5 mM; Sigma-Aldrich) to isolate monosynaptic responses. Inhibitory currents in some cases got pharmacologically blocked by bath application GABA_A_ antagonist Gabazine (10μM; Sigma-Aldrich). For pharmacological testing of Substance-P effect, Substance-P (1μM, Tocris) was bath applied in the aCSF. All parameters were analyzed by procedures custom-written in MATLAB (MathWorks, USA).

### Voltage imaging

After 4-5 weeks of Voltron-ST expression, mice were sacrificed and ex vivo brain slices of 250 μm were prepared. After ~30 min of incubation in 50 nM Janelia-Fluor 585 dye dissolved in aCSF, slices were transferred to the recording chamber of the electrophysiology setup. For the imaging we used a digital sCMOS camera (Orca Fusion-BT, Hamamatsu, Japan), frame triggers were sent to the camera with a Arduino Micro microcontroller (Arduino Uno) with 600 Hz. For all-optical imaging we used a dual-band excitation filter (ZET488/594, Chroma) to excite the JF-585 and deliver 473 nm light for optogenetic stimulation. The 585 nm light excitation intensity was ~10 mW/mm^2^ and 473 nm light intesity was ~2.5 mW/ mm^2^ at the slice plane and was delivered by Spectra X (Lumencore) LED light source. JF-585 fluorescent emission was collected with a 20X 1.0 NA water immersion objective (XLUMPLFLN20XW Plan Fluorit, Olympus). Emitted light was separated from the excitation light with a band-pass emmission filter (ET645/75, Chroma) and with a dichroic mirror (T612lprx, Chroma). Magnification was decreased with U-ECA magnification changer to 0.5x. To acquire videos we used the free LiveImage software triggered by the Arduino to synchronize acquisition with the frame triggers.

### Voltage-Seq

#### Neuronal soma harvesting

After voltage-imaging of each FOV and consequent on-site analysis in VoltView, we approached the suggested somas one-by-one with a harvesting pipette (1.8-2.5 MΩ) containing (in mM): 90 KCl, 20 MgCl2. The entire soma of the selected neuron was aspirated into the pipette within a few seconds by applying mild negative pressure (−50 mPa) measured with a manometer. Switching from DIC to the fluorescent optics to visualize the Voltron-expressing neurons during and after aspiration could confirm the succesful harvesting process. Next, the harvesting pipette was pulled out of the recording chamber and then with the micromanipulator, carefully navigated over and inside a 0.2 ml tube under visual guidance observed by a 4X Olympus air objective (Olympus, Tokyo, Japan). Applying positive pressure, the harvested neuron (~0.5 μl) was ejected into a 4 μl drop of lysis buffer consisting of 0.15% Triton X-100, 1 U/μl TaKaRa RNase inhibitor, 2.5 mM dNTP, 17.5 mM DTT and oligo dT primer (2.5 μM) pre-placed in the very tip of the 0.2 ml tight-lock tube (TubeOne). Through the 4X air objective we could observe the line of small air bubbles coming out of the harvesting pipette tip into the lysis buffer as a confirmation of the completed ejection. The resultant sample (~4,5 μl) was spun down (15-20 s), placed to dry ice and stored at −80 °C, later subjected to in-tube reverse transcription (RT).

#### Single-cell RNA sequencing

##### Library prep

Smart-Seq2 (SS2) libaries were prepared as previously described except for the following changed: SS2 cDNA was amplified for 22 cycles, cDNA 1ng was tagmented and amplified with custom 10bp indexes. Libraries were sequenced using a 150 cycle Nextseq 550 kit (paired end, 74bp reads).

##### Sequencing QC

Reads were aligned with the mm10 genome using zUMIs (version 2.9.5) using STAR (2.7.2a) with transcript annotations from GENCODE (version M25). Quality control reports for genes detected was performed with intron+exon alignments from zUMIs. Qualimaps was used to report the total reads aligned to intronic and exonic regions.

##### Differential expression and clustering

All expression analyses were performed with exonic reads only. The package Seurat in R was used to perform clustering analyses. Briefly, Smart-seq2 data was normalized to account for sequencing depth (gene count divided by total counts, multiplied by a scale factor of 10000 and log normalized) and the 1000 top variable features were found. Expression for each gene was scaled around 0 and a linear dimensionality reduction (PCA) was performed. For the clustering analysis, the K-nearest neighbor graph was constructed with the first 5 principal components and a Louvian algorithm iteratively grouped cells together using a resolution of 0.75 to obtain 2 clusters. Markers for each cluster were found using the Wilcoxon rank sum test. The heatmap for UMAP clustered cells and scaled marker expression was generated with ComplexHeatmap.

### Anatomical 3D mapping

Spatial mapping of imaged neurons was done using the common coordinate framework 4 (CCF4)^20^. X and Y coordinates were extracted from the videos relative to the top left corner and when Z coordinates were identified, the FOVs were mapped using „rails” generated with SHARPtrack (Extended Data Fig 5a). The package is originally used to track electrodes, but it is higly suitable for constructing reference points or lines in 3D and extract the coordinates to support 3D mapping. Our reference lines went all along the A-P axis of PAG on the most dorsal, lateral and ventral edges and within the PAG they were spaced by the distance of the size of FOVs recorded. “FOV fields” were registered during data acquisition and analyzed offline to calculate X,Y,Z coordinates of each imaged neuron.

#### Spatial core-density mapping

For each o-phys cluster, for each neuron, the number of neighbors in a 300 μm radius was calculated. The density core of a cluster was defined by the neuron with the highest number of neighbors. The spatial distance of the X,Y,Z coordinate of each neuron of the cluster was calculated from the X,Y,Z coordinate of the density core neuron. On the 3D core-density maps, neurons were color coded by the distance from the density core starting in red for closest towards blue with highest distance, transparency followed the color code with blue being invisible.

#### Axonal mapping

Each coronal cut PAG brain section was manually mapped to an AP coordinate (z) and x-y coordinates for each fluorescent image pixel (axonal fluorescent labeling) was segmented out in ImageJ. For mapping coronal slices of axonal masks we used the reference lines generated with SHARP-track and fitted the slices in the 3D PAG model with a custom-written script in MATLAB.

### Data analysis

#### VoltView analysis

The on-site and detailed analyis was written in MATLAB (MathWorks) using custom-written scripts. In major steps, the recorded videos from ‘.CXD’ files were imported to MATLAB using bioformats 6.11 package (OME - https://www.openmicroscopy.org/). Multiple sweeps were recorded in the same video in series. Putative lost frames were found and filled up. Camera artifact frames – marking the end of sweeps – were removed. Frames of the baseline before the optogenetic stimulation was averaged, frame average was local-equalized and contrast-enhanced specifically optimized with Voltron-JF585 imaging. These pre-processed images were used for the Cellpose segmentation of somatic ROIs. After the detection of ROI contours we designated the pixels of concentrical ROIs (Inner and Outer) around the somatic ROI. The number of pixels of Inner and Outer were scaled to be within ±5% of the somatic ROI surface so that signal-to-noise can be compared. We extracted the optical physiology (o-phys) traces from all somatic, Inner and Outer ROIs. We calculated the dF/F from the o-pys traces and used exponential fitting to correct for bleaching. Basic features, o-AP peaks, subthreshold kinetics (o-Sub) and Burst activity was detected first. O-AP detection used the noise of the baseline for each ROI to calculate a cutoff threshod at 2.5-times the std – this may differ based on the illumination light intensity of voltage imaging as the signal-to-noise ratio is affected by that. We used polynomial fitting for straightening the dF/F traces before the o-AP detection (which we also validated by data from simultaneous whole-cell patch-clamp recordings). O-sub was the moving average of the 600 Hz o-phys trace by default. As wider o-APs – based on o-AP-half-width measurement - needed different moving average, we scaled it accordingly for o-Sub detection. Burst detection used a slower moving average with averaging 100 points (o-Sub-slow) against the o-Sub, where the o-Sub went above the o-Sub-slow, the periods were considered as putative burst periods. Next, o-AP detection was overlayed and compared to putative burst periods and if multiple o-APs were on these fast depolarizations, they were considered to be bursts (this was optimized and validated with simultaneous whole-cell patch-clamp and all-optical voltage imaging as in Extended Data Fig 3c-e). Using the basic features (o-AP peaks, o-Sub, Burst) we compared the 3 o-phys traces of the ROIs (somatic, Inner, Outer) with the amplitude of o-APs, the amplitude of o-Sub and the length and number of bursts across the traces. If the amplitudes and burst length were decreasing going from the somatic ROI towards the Inner and towards the Outer zones, the ROI was comfirmed to be the source of the response. In case if the signal parameters became stronger from the soma towards the Outer zone, the ROI was discarded. Traces were denoised using wavelet transformation or with multiscale local 1-D polynomial transformation. Optical blue stimulation artifacts were removed. As the Voltron-imaging with our analysis could detect overall depolarization and hyperpolarization compared to baseline level during Op, oEPSPs and o-IPSPs were detected using the o-Sub thresholded with ±5 x sd of the baseline and searching changes in the first derivative which represent inreases of o-EPSPs or decreases of o-IPSPs. All the calculated values and features were stored in struct variables so that VoltView can call them for the ROI explorer plotting and for the on-site analysis.

#### Classification and clustering of o-phys

Parameters for hierarchical clustering were extracted from the o-phys traces by custom-written routines in MATLAB (MathWorks). Parameters were averages of 6-7 sweeps recorded from each ROI. “o-EPSP” and “o-IPSP” was the number of o-PSPs during the 20 Hz Op. “Paired Pulse 2-1” compared the second o-PSP to the first o-PSP amplitude. “Paired Pulse 3-2” compared the third o-PSP to the second o-PSP amplitude. “Subthr Slope q2-q1” compared the mean amplitude of o-Sub between second 250 ms and the first 250 ms. “Subthr Slope q4-q1” compared the mean amplitude of o-Sub between last 250 ms and the first 250 ms. “AP onset” is the delay of the first o-AP during 20 Hz Op. “AP bimodality coeff” calculated Sarle’s bimodality coefficient as the square of skewness divided by the kurtosis, value for uniform distribution is 5/9, values greater than that indicate bimodal (or multimodal) distribution. “AP bimodality binary” gave 1 for bimodal (when coefficient was above 5/9) and 0 for non-bimodal o-AP peak distribution. “AP % in burst” quantified the number of o-APs inside burst periods. “Burst AP freq (Op)” quantified the firing rate inside detected bursts during 20 Hz Op. “AP freq 2nd/1st (Op)” quantified the change of firing rate during the 20 Hz Op comparing the second half (Op-2) with the first half (Op-1). “Burst AP freq (Post)” quantified the firing rate inside detected bursts after 20 Hz Op. “AP freq 2nd/1st (Post)” quantified the change of firing rate after the 20 Hz Op comparing the second half (Post-2) with the first half (Post-1). “AP num” gave the total number of o-APs detected on the baseline of all 6-7 sweeps. “AP freq (Baseline)” was the average firing rate on the baseline before the 20 Hz Op. “AP freq (Op-1)” was the average firing rate on the first half of 20 Hz Op. “AP freq (Op-2)” was the average firing rate on the second half of 20 Hz Op. “AP freq (Post-1)” was the average firing rate on the first half of Post, after 20 Hz Op. “AP freq (Post-2)” was the average firing rate on the second half of Post, after the 20 Hz Op. “Burst num (Op-1)” was the number of burst periods during the first half of 20 Hz Op. “Burst num (Op-2)” was the number of burst periods during the second half of 20 Hz Op. “Burst num (Post-1)” was the number of burst periods during the first half of Post, after the 20 Hz Op. “Burst num (Post-2)” was the number of burst periods during the second half of Post, after the 20 Hz Op. “Burst length (Op-1)” was the average length of burst periods during the first half of 20 Hz Op. “Burst length (Op-2)” was the average length of burst periods during the second half of 20 Hz Op. “Burst length (Op-2/Op-1)” was the change of burst length from Op-1 to Op-2. “Burst length (Post-1)” was the average length of burst periods during the first half of Post, after the 20 Hz Op. “Burst length (Post-2)” was the average length of burst periods during the second half of Post, after the 20 Hz Op. “Burst length (Post-2/Post-1)” was the change of burst length from Post-1 to Post-2. Agglomerative clustering was done by Ward’s algoritm, Euclidean distance for the cutoff and for definition of the number of clusters was chosen based on the slope-change of number of clusters vs the Euclidean distance-cutoff.

#### VoltView classifier

We used the unbiased hierarchical clustering of the connectome dataset and calculated the cluster centroids for each o-phys cluster. We had all the 29 parameters in the cluster centroid. We used two parallel classifications and used their consensus. First, for each neuron we calculated the square root of sum squared differences compared to all the cluster centroids and ranked the distances to choose the closest 3 clusters for each neuron. Next for each neuron we calculated the level of correlation to all the cluster centroids, ranked the correlation and choose the 3 highest correlating clusters. If the consensus of the two parallel classifications agreed on one or more clusters, we compared the ranks and assigned the closest cluster to each neuron.

#### Spatial cluster density

The observed minimal distances in space from a reference cell to a cell in the target cluster were calculated, as well as the expected minimal distances by chance, based on 1000 shuffles of cluster identity, for all cells. The distance metric was calculated for each cell by:

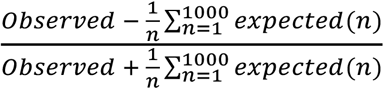

and is in range −1:1, where negative values indicated that the minimal distance of the reference cell to a cell in the target cluster is smaller than by chance and positive values indicated greater distance than by chance.

---

**Extended Data Fig.1:**
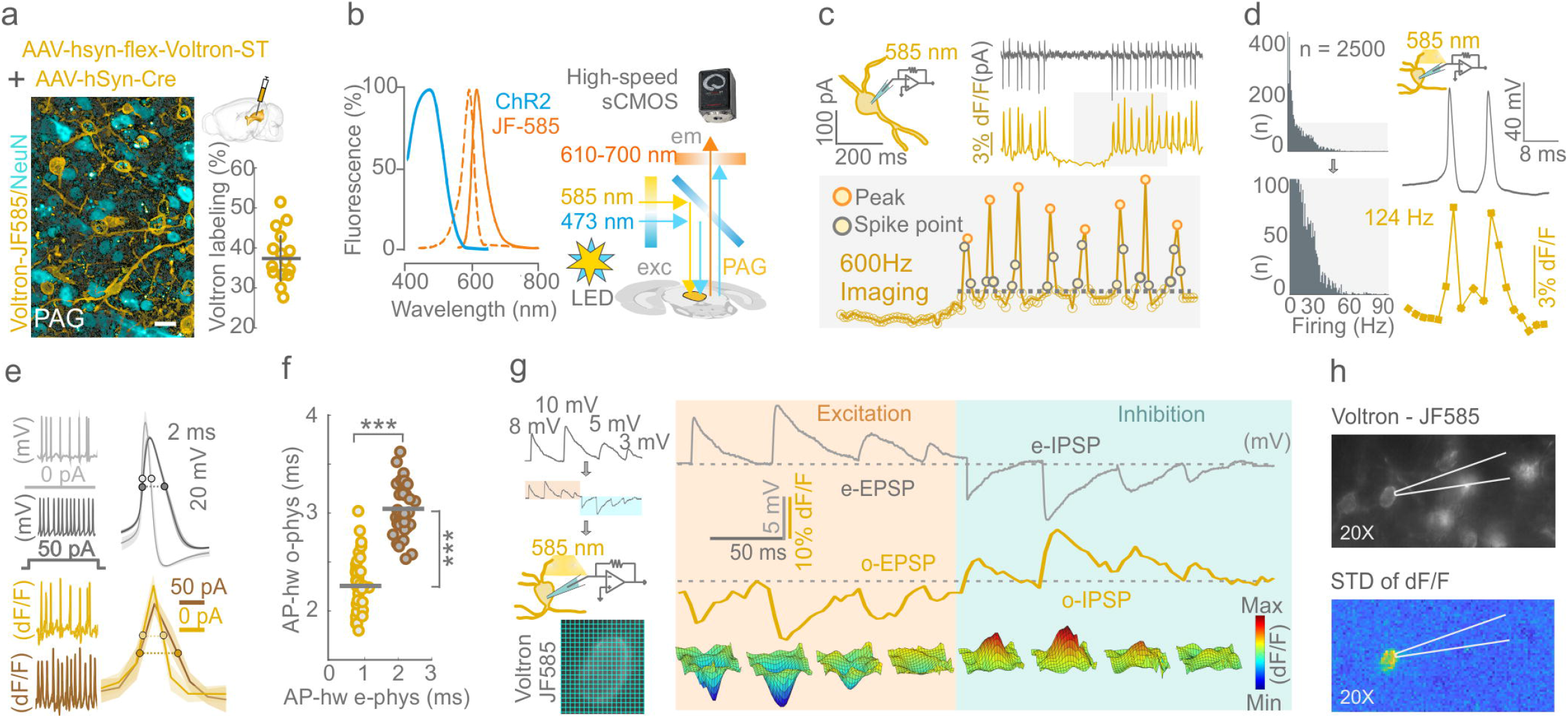
Optical setup, frame rate and detection limits of Voltron imaging. All Voltron traces were reversed except in panel g. **a,** NeuN staining and Voltron labeling with JF-585 (left, Scale bar is 20 μm). Scheme of Voltron-expressing AAV injection to PAG, hive plot (mean±sd) of the percentage of JF-585-Voltron-ST neurons/total number of neurons in randomly designated areas of PAG. **b,** Excitation, and emission profile of ChR2 (blue) and JF-585 (orange) (left). Optical setup of all-optical voltage imaging *ex vivo* (right). **c,** Scheme of simultaneous cell-attached patch-clamp recording and voltage imaging (top left), APs from a PAG neuron with time-aligned traces of e-phys (black) and o-phys (gold) from the same neuron (top right). Insert of o-phys trace acquired at 600 Hz framerate with AP peaks (Peak points) and points between AP-threshold and peak (Spike points), dashed line marks AP-threshold (bottom). **d,** Histogram of detected firing rate in 2500 Pag neurons; insert highlights that in our dataset during 20 Hz Op we detected firing rates up to 60Hz (left), we validated the highest firing rate we could detect is ~124 Hz (right). **e,** E-phys trace at 0 pA (light gray) and 50 pA (dark gray) depolarizing current injection with the corresponding o-phys traces at 0 pA (gold) and 50 pA (brown) (left). AP shape of e-phys (light gray, dark gray) and o-phys spikes (gold, brown) at 0 pA and 50 pA current injection (right). **f,** Half-width of APs compared across 0 pA and 50 pA current injection of e-phys (p = 1,9e-49) and o-phys (p = 6,4e-16) recordings (***p<0.001). **g,** Scheme of symmetric voltage command to test the subthreshold sensitivity of the Voltron sensor (top left). The applied voltage command had amplitudes of 8, 10, 5, 3 mV and −8, −10, −5, −3 mV. Scheme of simultaneously voltage-imaged and whole-cell recorded neuron (middle left). Soma of a JF-585-Voltron-ST neuron, representing the pixel-binning for the surface plots (bottom left). Application of the symmetric voltage command in whole-cell voltage-clamp during voltage imaging, e-phys recording (top, black), with the corresponding o-phys (bottom, gold) trace from the same recorded neuron (avg. of 10 sweeps). Surface plots of dF/F of the recorded soma with 4 by 4 binned pixels (bottom). **h,** JF-585-Voltron-ST neurons during imaging and whole-cell patching in 20x magnification (top). Std of dF/F of the whole-cell-recorded neuron, with the surrounding neurons not depolarized (bottom).

**Extended Data Fig.2.:**
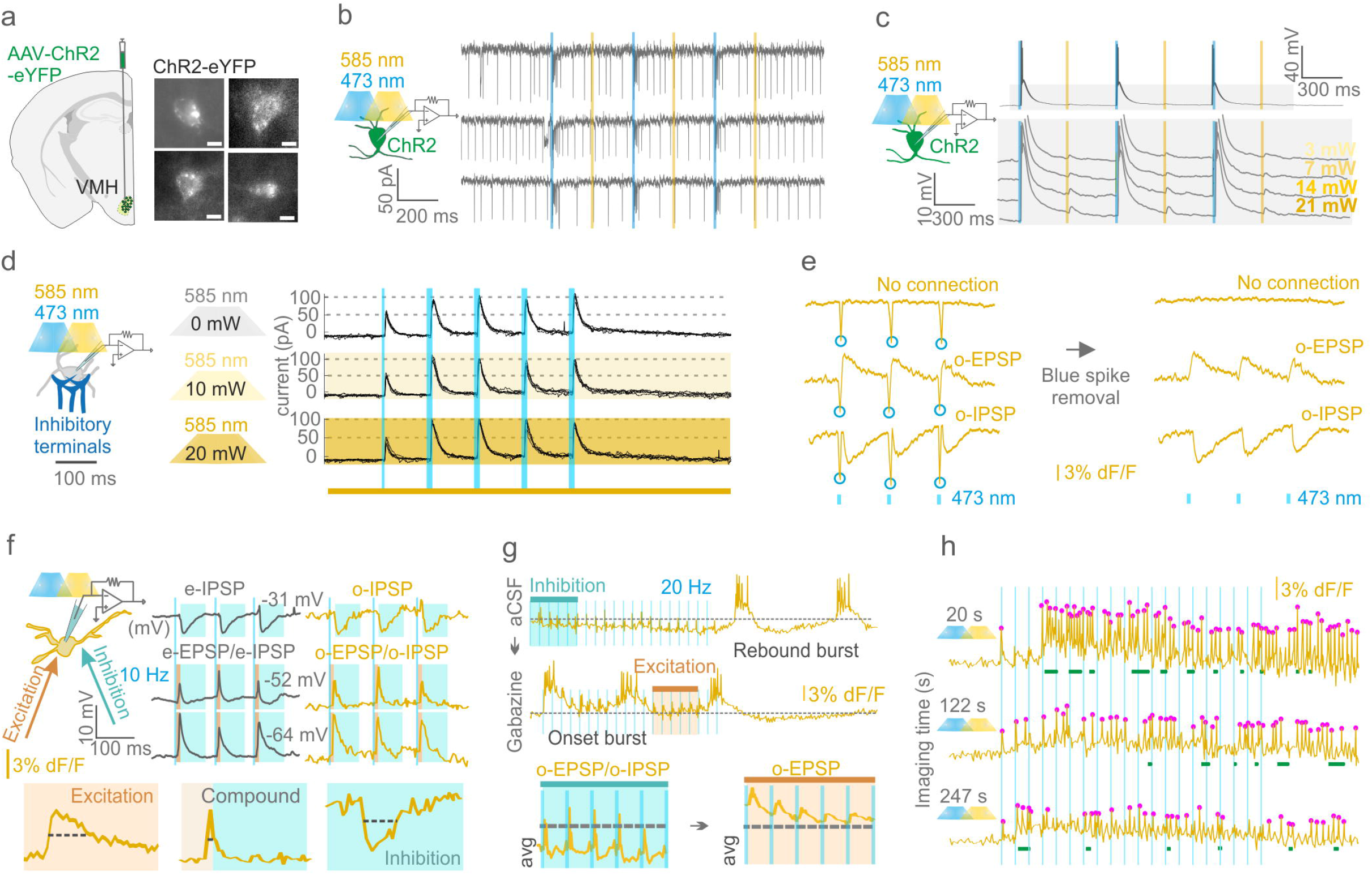
Testing Voltron imaging and validation of all-optical recordings. **a,** Scheme of ChR2 expression in VMH with ChR2 expressing somas (left, scale bar 10 μm). **b,** Scheme of the experiment (left), stimulating VMH neuron with 473 nm and 585 nm light during cell-attached recording (right). Illumination with 473 nm was able to activate ChR2, but 585 nm light did not evoke a response. **c,** Scheme of experiment (left), ChR2-expressing VMH neuron during whole-cell patch-clamp recording and optogenetic stimulation with 473 nm and 585 nm light (right). Gray area enlarged below, 3mW 473 nm light could activate ChR2 and caused strong depolarization, the same light power at 585 nm only evoked a small (~1 mV) voltage change. Recordings from top to bottom were stimulated with 3,7,14, 21 mW 585 nm light. The maximum 21 mW light power with 585 nm evoked ~2-3 mV depolarization in the recorded neuron. **d,** Scheme of the experiment (left). Right, whole-cell voltage-clamp recordings. We induced incomplete synaptic release with the first narrower (0.75 ms) 473 nm light stimulation and maximal release with four wider (3 ms) 473 nm stimulations. 585 nm light with 0 mW, 10 mW and 20 mW illuminated the recorded neuron during repetition of the incomplete release Op protocol. We confirmed that synaptic ChR2 illuminated by 20 mW 585 nm light did not activate to result in a maximal transmitter release. **e,** Each 473 nm light pulse during 585 nm illumination of the Voltron sensor induced a narrow “blue spike” artefact. These serve as time stamps of optogenetic stimulation and are reversed in polarity to o-APs, analysis removes them from all Voltron recordings. **f,** Left, schematic illustration of a PAG neuron receiving both excitatory and inhibitory synaptic inputs during Op. Side by side e-phys (average of 5 traces, black) and o-phys (average of 5 traces, gold) traces at different command voltages (−31 mV, −52 mV, −64 mV) with inhibitory input dominating (e-IPSP, o-IPSP) at most depolarized (−31 mV) and excitatory input dominating (e-EPSP, o-EPSP) at most hyperpolarized (−64 mV) command voltages. Compound PSP half-width is extremely narrow compared to the ones during excitation (in brown) or inhibition (in cyan). **g,** All-optical traces of a PAG neuron showed inhibition (insert, compound o-PSPs, cyan) during Op of VMH input in aCSF and exerted rebound burst firing after the Op (Rebound burst); same PAG neuron in Gabazine (10 μM) showed elimination of the GABA_A_-mediated inhibition resulting in excitation (insert, o-EPSPs, brown) onset burst firing during Op (Onset burst) **i,** 3 sweeps from the same neuron during repeated imaging after 20, 122, 247 s imaging time demonstrates bleaching, which became robust after 3-4 minutes of imaging.

**Extended Data Fig.3:**
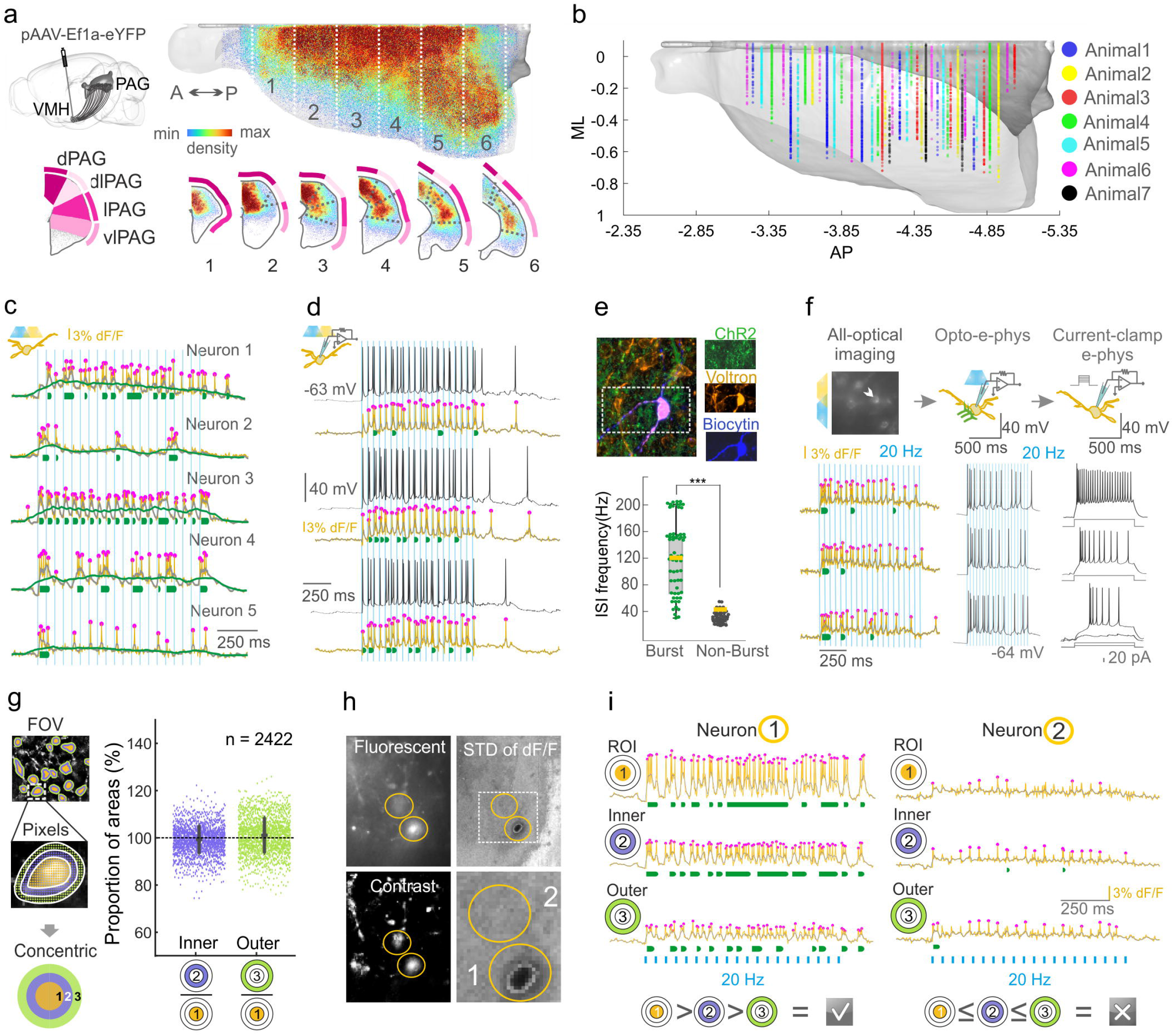
Axonal map, Imaged coordinates, Burst detection, Concentric. All Voltron traces were reversed. **a,** Expression of eYFP in the VMH-PAG pathway (top left); PAG hemisphere with 3D reconstruction of VMH axonal density in the PAG, color codes density from blue (low) to red (high) (top). Coronal scheme shows the color code of PAG subregions (bottom left); Coronal bins (1: −3,25/-3,65 2: −3,65/-3,95; 3: −3,95/-4,25; 4: −4,25/-4,55, 5: −4,55/-4,85, 6: −4,85/-5,15 mm to Bregma) show distribution of VMH axons across PAG subregions (bottom). **b,** A-P coordinates of imaging planes with 6911 imaged neurons (Dots in the planes are individual neurons (6911 neurons/7 animals). **c,** All-optical-imaged neurons (Cell1-Cell5) with different burst types. Magenta dots mark the detected o-APs on o-phys traces (gold), O-sub is the moving average of 10 points (gray), the moving average of 100 points (green). Green lines under the o-phys traces indicate the detected bursts. **d,** PAG neuron during all-optical imaging and whole-cell patch-clamp recording. Corresponding o-phys (gold) and e-phys (black) sweeps, magenta dots mark the detected o-APs, green lines under the o-phys traces indicate the detected bursts. **e,** Biocytin-filled JF-585-Voltron-ST PAG neuron between the ChR2-expressing VMH axonal terminals (top). Inserts from top to bottom show ChR2 expression, Voltron expression and biocytin labeling (top right). Box plot compares the frequency of ISIs where burst was detected (green dots) compared to ISIs where bursts were not detected (gray dots). Dots represent individual ISIs from 6 o-phys sweeps of the shown biocytin labeled neuron (mean±sd t-test, ***p<0.001) (bottom). **f,** FOV of JF-585-Voltron-ST PAG neurons (top), white arrow points at the neuron chosen for whole-cell patch-clamp recording (top left, scale bar: 20 μm)) based on the detection of onset-bursts (green lines) on its o-phys traces (gold) (bottom left). Scheme of the recording (top middle), 3 e-phys traces, 20 Hz Op during whole-cell patch clamp recording confirmed the onset-bursting predicted by the o-phys recording (bottom middle). Scheme with the same JF-585-Voltron-ST neuron during whole-cell current-clamp (top right), intrinsic excitability characterization confirmed a burst firing electrophysiological type validated the prediction of o-phys detection of intrinsic features (bottom right). **g,** Concentric analysis generates 2 concentric areas (2: purple (Inner), 3: green (Outer)) around somatic ROIs (1: gold) (left). The number of pixels is scaled to make 1,2 and 3 similar in surface for side-by-side comparison of signal to noise; scatter plot of surface ratio between Inner/Roi (sd = ±5.2%) and Outer/Inner (sd = ±8.3%) areas of n = 2422 neurons. **h,** Fluorescent image of 2 JF-585-Voltron-ST neurons (yellow circles) (top left), same neurons on a local equalized and contrast enhanced image better visualizes the somatic JF-585 signals (yellow circles) (bottom left), ±sd of dF/F voltage signal of the same neurons (yellow circles), insert shows that in contrast to Neuron1, Neuron2 did not have activity during all-optical imaging. **i,** Concentric analysis of the 2 neurons from h. Neuron1 displayed strong firing and bursting activity (ROI), which could be detected in the concentric zones (Inner, Outer) with decreased o-AP amplitude, less and shorter burst detections (left). Neuron2 displayed o-APs in its soma (ROI), but compared to concentric zones (Inner, Outer) the number and amplitude of o-APs increased indicating that the source of the signal is not Neuron2. The Outer of Neuron1 and the Outer of Neuron2 showed similarities indicating the signal source of Neuron2 was probably Neuron1.

**Extended Data Fig. 4:**
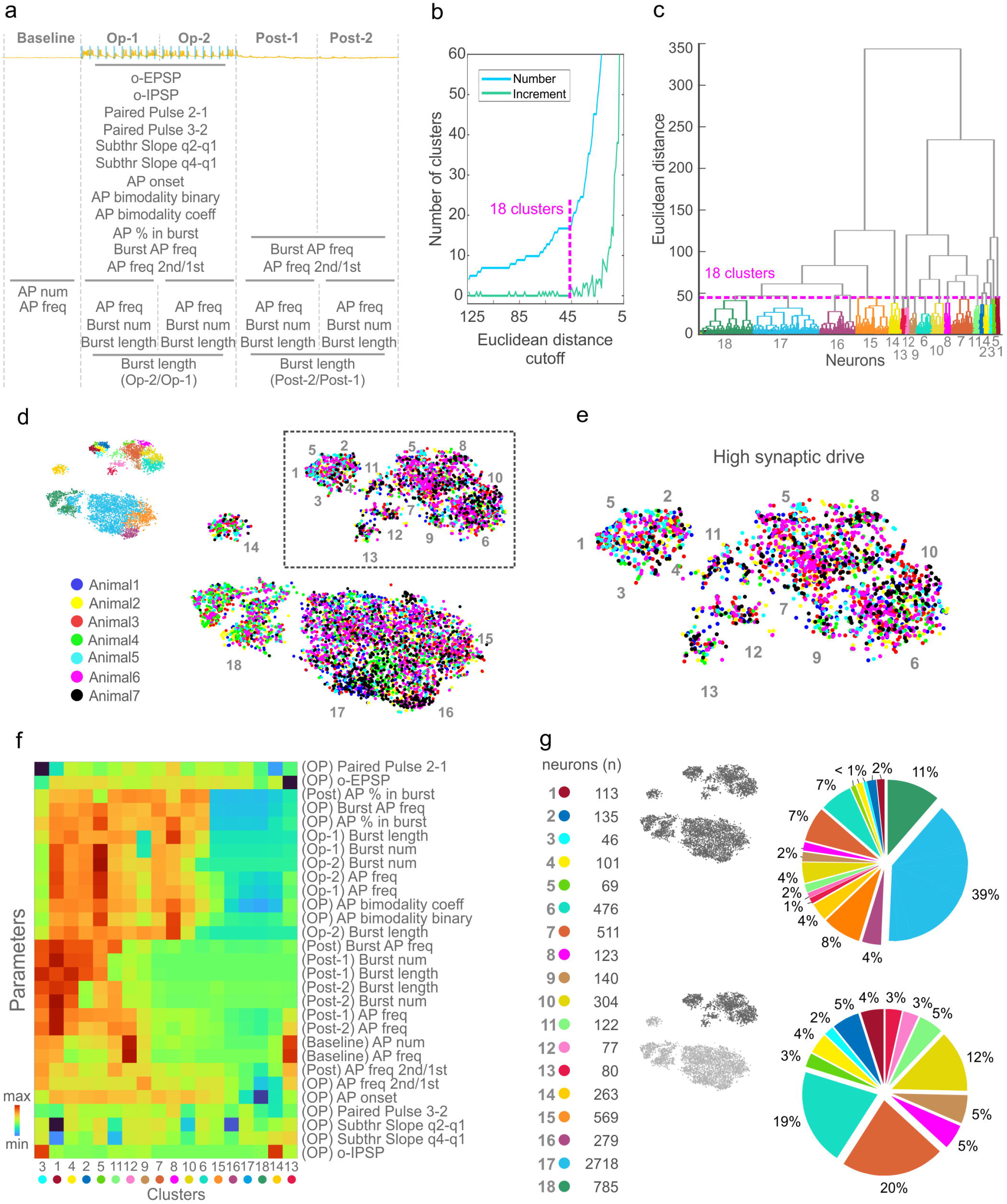
Clustering of VMH-PAG o-phys PRTs. **a,** 29 parameters extracted from the o-phys traces used for the hierarchical clustering of o-phys types. Scheme on top details the temporal segments of all-optical sweeps with Op-1: first half of Op, Op-2: second half of Op, Post-1: first half of after-Op, Post-2: second half of after-Op. **b,** line plot of the relation of number of clusters versus Euclidean distance cutoff of the agglomerative hierarchical clustering (blue) and the first derivative of the same curve showing the increase in the number of clusters versus Euclidean distance cutoff of the agglomerative hierarchical clustering (green). **c,** Hierarchical dendrogram of o-phys of the 6911 neurons with the Euclidean cutoff resulting in 18 clusters (color code matches Fig. 2f,g). **d,** tSNE plot of VMH-PAG o-phys clusters with color coding the imaged neurons by animal (1-7). **e,** High synaptic drive clusters enlarged from d, to show the lack of batch effect across tSNE regions and clusters. **f,** Heat map of clustering parameters versus the 18 o-phys clusters show characteristics of the distinct o-phys clusters. **g,** Quantification of neurons in each o-phys cluster (left). Pie chart of cluster proportions (%) in all the VMH-PAG connectome (top right), and in the high synaptic drive clusters (bottom right).

**Extended Data Fig. 5:**
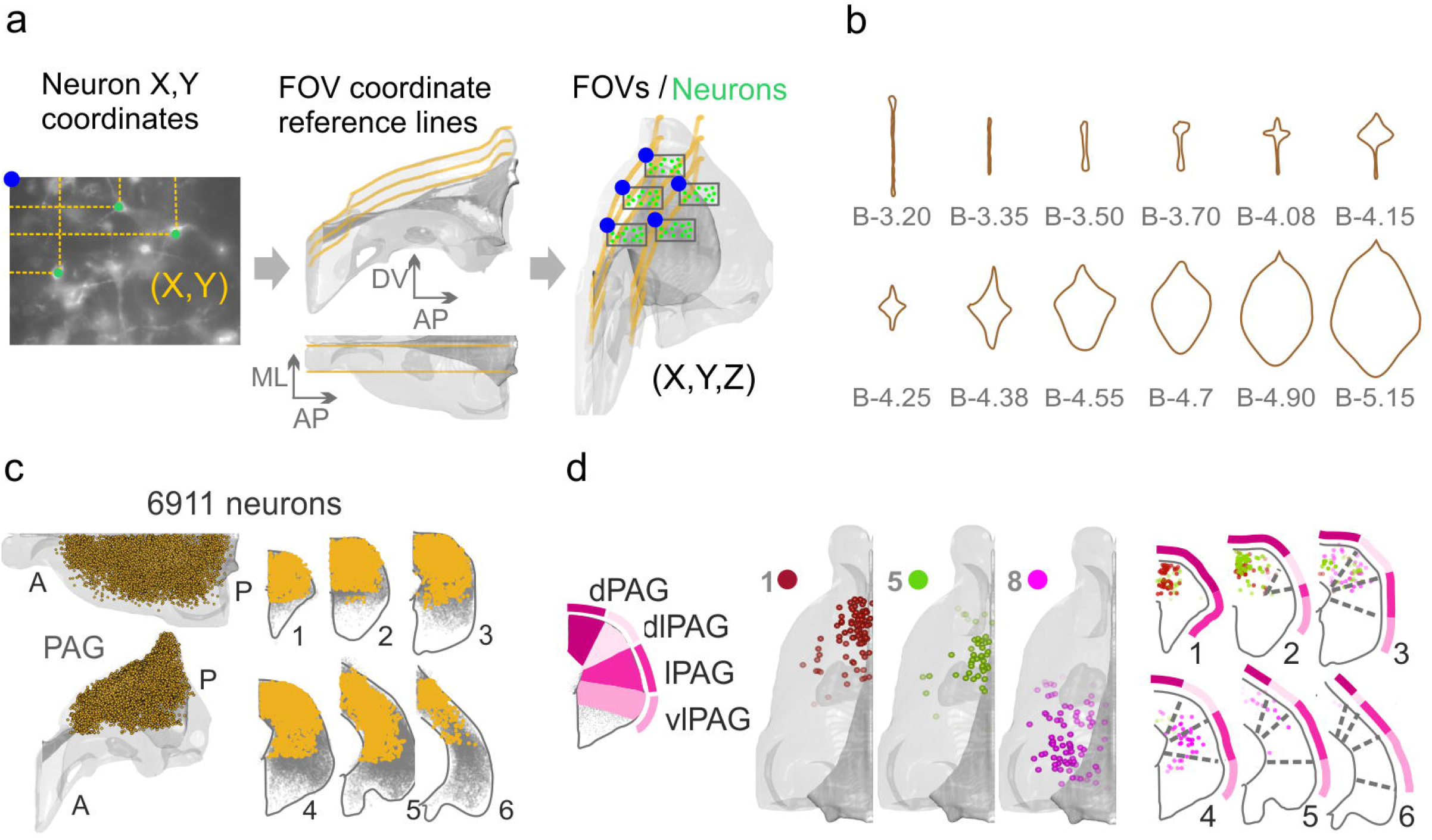
Spatial mapping of all-optical connectome. **a,** 3D mapping of the all-optical voltage-imaged PAG neurons; FOV with JF-585-Voltron-ST neurons illustrate the extraction of X and Y coordinates relative to the top left corner of each FOV (blue) (left), reference lines were spaced away by the size of the FOVs (middle) and was used to rail the FOVs upon mapping up to the identified A-P coordinate (right). **b,** Line drawings of the aqueduct (AQ) along the A-P axis of PAG, templated with coronal PAG sections of the Allen reference atlas. A-P coordinate was defined based on the shape of the AQ **c,** 3D hemisphere of PAG with 6911 neurons (gold) dorsal view (top), transverse view (bottom) (left), Coronal bins (1: −3,25/-3,65 2: −3,65/-3,95; 3: −3,95/-4,25; 4: −4,25/-4,55, 5: −4,55/-4,85, 6: −4,85/-5,15 mm to Bregma) show the coverage of PAG subregions (right). **d,** 3D PAG model of one hemisphere for v-phys cluster 1,5,8 with the density-core mapping of postsynaptic neurons colored by cluster identity (left), merging the spatial coordinates of cluster 1,5 and 8 on the same coronal bins as in c (right).

**Extended Data Fig.6.:**
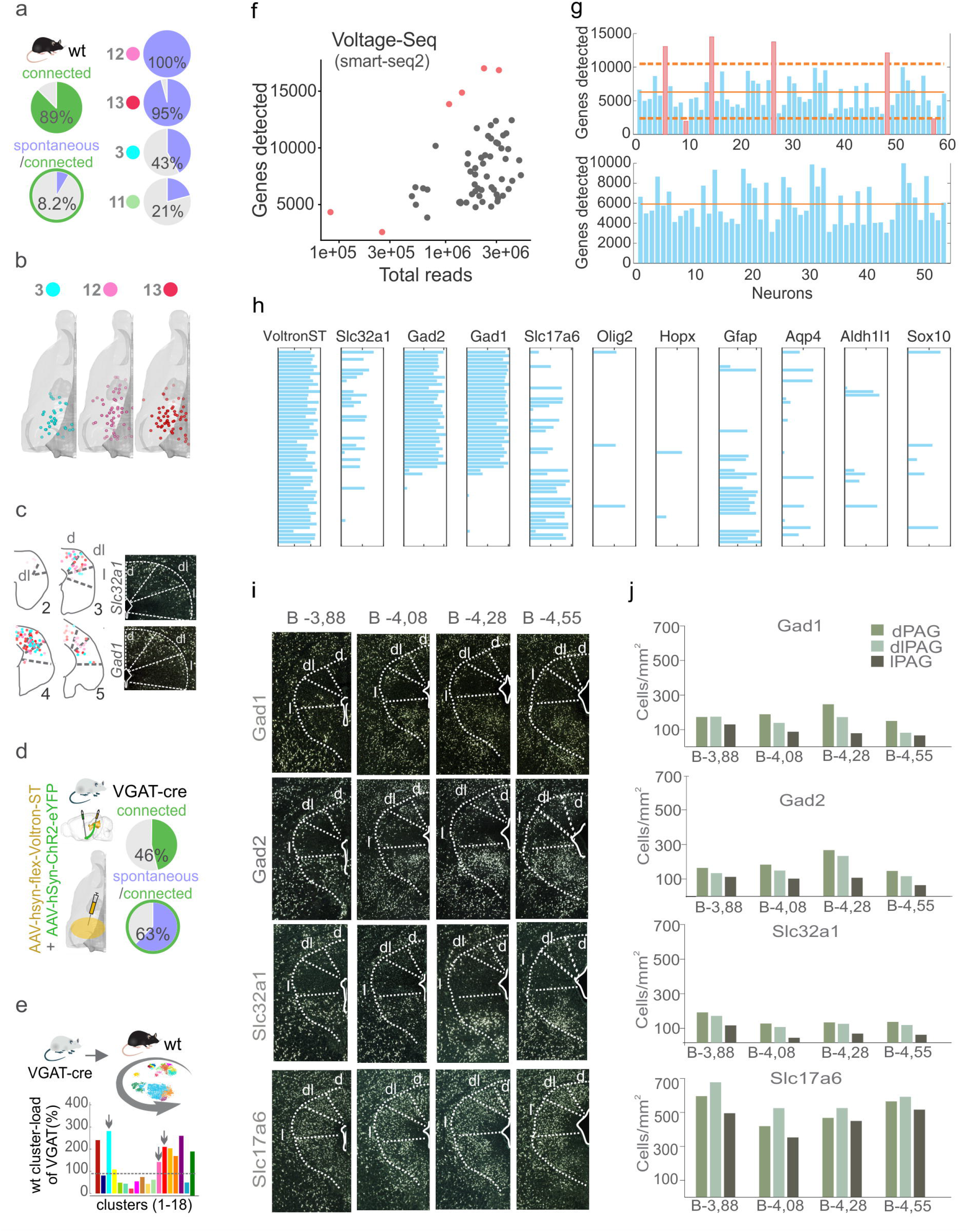
Spontaneously active clusters, scRNA-seq, ISH quantification. **a,** Pie chart of detected connections in VMH-PAG connectome of wt mouse (top left), and pie chart of spontaneously firing postsynaptic neurons within connected neurons (bottom left). Pie charts of the ratio of spontaneously active neurons within cluster 12, 13, 3, 11 (right). **b,** 3D PAG model of one hemisphere for o-phys cluster 3,12,13 with the density-core mapping of postsynaptic neurons of the three clusters color coded by o-phys cluster identity. **c,** Coronal bins of PAG (2: 3,65/3,95; 3:3,95/4,25; 4: 4,25/4,55, 5: 4,55/4,85) merging the spatial coordinates of cluster 3,12,13 (left), spatial distribution of *Slc32a1* and *Gad1* ISH signal (modified from the Allen ISH database) (right). **d,** Scheme of ChR2 and Voltron-ST expression in the posterior portion of PAG of VGAT-Cre mice (left), pie chart of detected connections in our VGAT-cre VMH-PAG connectome (top right), and pie chart of spontaneously firing postsynaptic neurons within the inhibitory VMH-PAG connectome (bottom right). Spontaneous firing compared to wild-type (8.2%) in (a) was 8-times higher (63.7%) in the VGAT data **e,** VGAT PRTs were classified by the classifier in VoltView to probe cluster-load of wild-type o-phys clusters with VGAT o-phys. Cluster-load of wild-tape clusters was divided by the cluster-load of VGAT o-phys to compare the proportional difference. Cluster 1,3,12,13 and subthreshold clusters (14,15,16,18) had the highest relative cluster-load with VGAT o-phys. Cluster1 was anatomically separated, and subthreshold clusters did not exert spontaneous firing; we chose cluster 3,12 and 13 for Voltage-Seq. **f,** Scatter plot of detected genes per number of reads with Smart-Seq2 library preparation protocol in Voltage-Seq data. **g,** Bar plot of number of detected genes in each 60 Voltage-Seq neurons (blue), mean (solid orange) and ±1.5 x sd (dashed orange), 6 discarded neurons with red bars (top); Bar plot of number of detected genes in quality controlled 54 Voltage-Seq neurons (blue) with new mean (solid orange) (bottom). **h,** Bar plot of Voltage-Seq RNA-transcriptome with the expression of *Voltron-ST*, *Slc32a1*, *Gad2*, *Gad1*, *Slc17a6 and glial markers Olig2*, *Hopx*, *Gfap*, *Agp4*, *Aldh1l1*, *Sox10*. Minor ticks are 5 and 10 TPM from left to right for each gene. **i,** ISH of PAG coronal slices (−3.88, −4.08, −4.28, −4.55 from Bregma) implemented from the Allen ISH database with inverted colors. GABAergic markers: *Gad1* (GAD67), *Gad2* (GAD65), *Slc32a1* (VGAT), and glutamatergic marker: *Slc17a6* (VGLUT2) shows the spatial distribution of GABAergic and glutamatergic neurons, respectively (top to bottom). **j,** Bar plots of ISH quantification (positive nuclei/mm^2^) in dPAG, dlPAG and lPAG in four A-P coordinates (−3.88, −4.08, −4.28, −4.55 from Bregma) for *Gad1*, *Gad2*, *Slc32a1*, and *Slc17a6* (top to bottom). This quantification was used to calculate the chance-level of finding GABAergic neurons in the PAG.

**Extended Data Figure 7:**
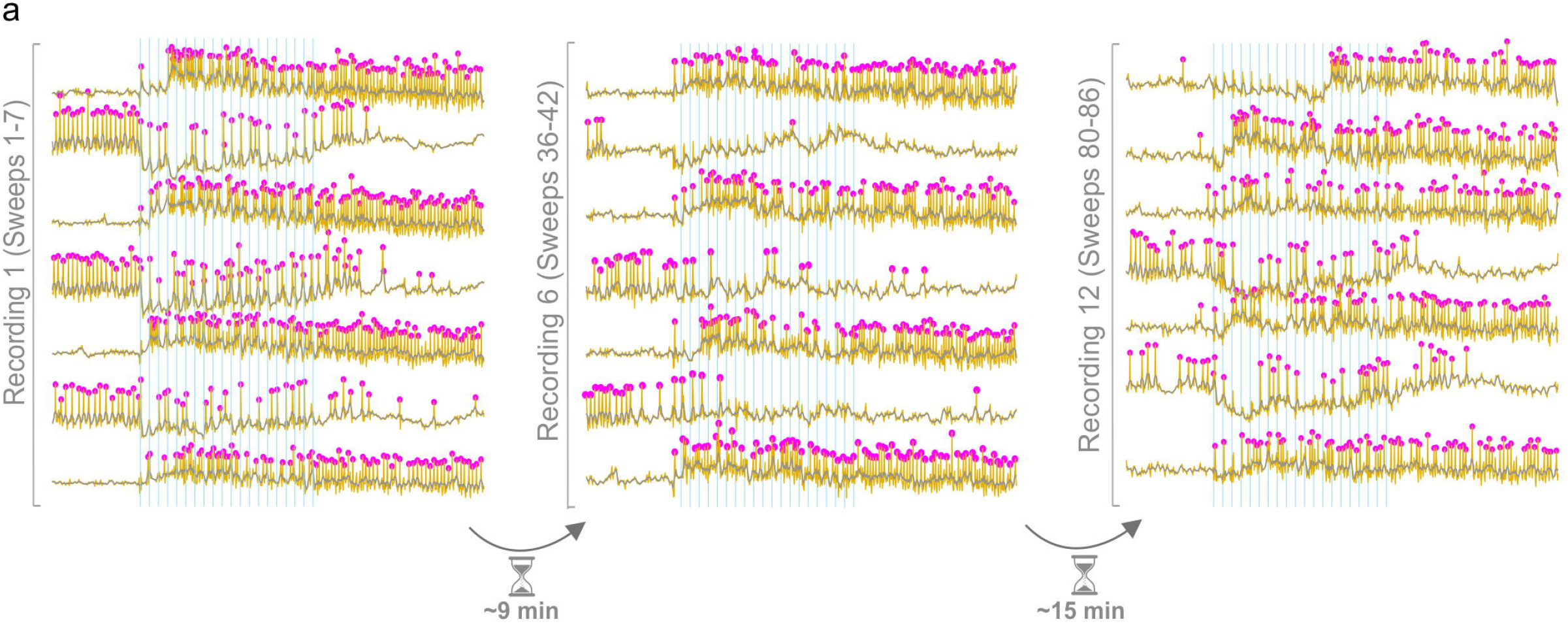
Stability of the Switch response. **a,** Example o-phys traces of the same Switch responder over time. First all-optical recording (left), all-optical recording 9 min later (middle), and all-optical recording 15 min after the second recording (26 min in the recording chamber). Switch behavior could be observed throughout more than 80 sweeps with a stable switching using our recording conditions, this validated the stability of Switch response and made it compatible with further detailed investigation for example in plasticity protocols or pharmacology experiments.

**Extended Data Figure 8:**
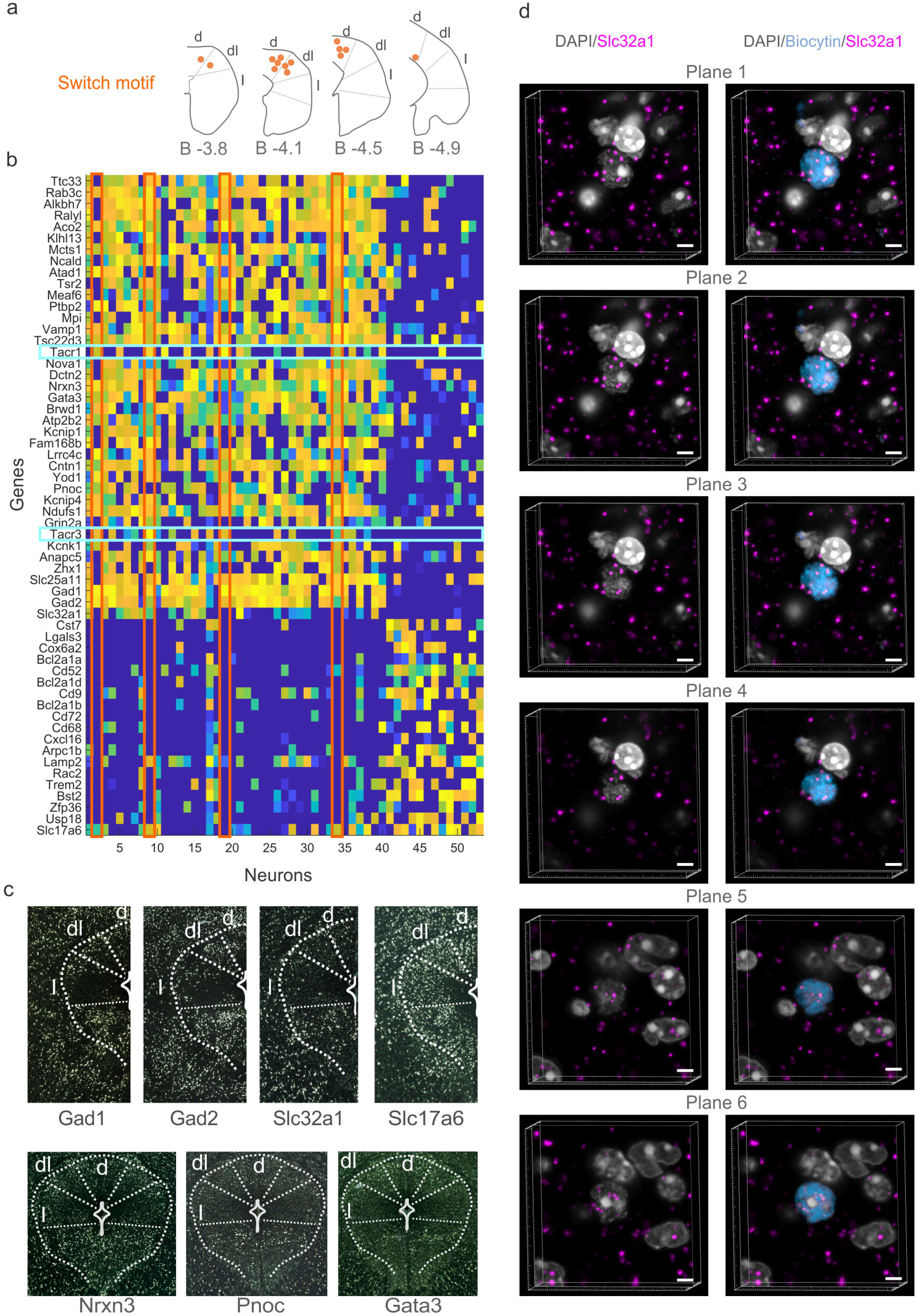
Differential expression, GABAergic markers, ISH. **a,** Anatomical position of Switch responders (orange dots) on coronal slices in d-,dl-,lPAG. **b,** Heatmap of DE genes; Switch responders (orange rectangles), express *Tacr1* and *Tacr3* (cyan rectangles). **c,** ISH of PAG coronal slices at B-4.08 implemented from the Allen ISH database with inverted colors. *Gad1*, *Gad2*, *Slc32a1*, and *Slc17a6* shows the spatial distribution of GABAergic and glutamatergic neurons, respectively (top); ISH of PAG coronal slices with DE-identified putative GABAergic markers (*Nrxn3*, *Pnoc*, *Gata3*) shows similar spatial distribution to that of GABAergic neurons. **d,** 6 differnet planes of *Slc32a1* (*magenta*) ISH on a biocytin-filled burst firing Switch responder shows nuclear colocalization of *Slc32a1*.

